# The FUR-like regulators PerRA and PerRB integrate a complex regulatory network that promotes mammalian host-adaptation and virulence of *Leptospira interrogans*

**DOI:** 10.1101/2020.11.05.369660

**Authors:** André A. Grassmann, Crispin Zavala-Alvarado, Everton B. Bettin, Mathieu Picardeau, Nadia Benaroudj, Melissa J. Caimano

## Abstract

*Leptospira interrogans*, the causative agent of most cases of human leptospirosis, must respond to myriad environmental signals during its free-living and pathogenic lifestyles. Previously, we compared *L. interrogans* cultivated *in vitro* and *in vivo* using a dialysis membrane chamber (DMC) peritoneal implant model. From these studies emerged the importance of genes encoding the Peroxide responsive regulators PerRA and PerRB. First described in in *Bacillus subtilis*, PerRs are widespread in Gram-negative and -positive bacteria, where regulate the expression of gene products involved in detoxification of reactive oxygen species and virulence. Using *perRA* and *perRB* single and double mutants, we establish that *L. interrogans* requires at least one functional PerR for infectivity and renal colonization in a reservoir host. Our finding that the *perRA*/*B* double mutant survives at wild-type levels in DMCs is noteworthy as it demonstrates that the loss of virulence is not due to a metabolic lesion (*i.e.*, metal starvation) but instead reflects dysregulation of virulence-related gene products. Comparative RNA-Seq analyses of *perRA*, *perRB* and *perRA/B* mutants cultivated within DMCs identified 106 genes that are dysregulated in the double mutant, including *ligA, ligB* and *lvrA/B* sensory histidine kinases. Decreased expression of LigA and LigB in the *perRA*/*B* mutant was not due to loss of LvrAB signaling. The majority of genes in the *perRA* and *perRB* single and double mutant DMC regulons were differentially expressed only *in vivo*, highlighting the importance of host signals for regulating gene expression in *L. interrogans*. Importantly, the PerRA, PerRB and PerRA/B DMC regulons each contain multiple genes related to environmental sensing and/or transcriptional regulation. Collectively, our data suggest that PerRA and PerRB are part of a complex regulatory network that promotes host adaptation by *L. interrogans* within mammals.

**Author Summary:** Leptospirosis is a neglected tropical disease with a worldwide distribution. Globally, ~1 million cases and ~60,000 deaths are reported each year. The majority of cases of human leptospirosis are associated with *Leptospira interrogans*. Infection begins when a naïve reservoir (or incidental) host comes into direct or indirect contact with urine from an infected reservoir host. While infection in reservoir hosts, including rats and mice, is generally asymptomatic, incidental hosts, including humans, may develop clinical symptoms ranging from mild flu-like illness to fulminant disease. The gene products required by leptospires for infection remain poorly understood. Herein, we establish that the FUR family regulators PerRA and PerRB function in parallel, contributing to infectivity and renal colonization in mice. By comparative transcriptomics, we identified >100 genes that were dysregulated in the *perRA/B* double mutant cultivated in rat peritoneal cavities, including the virulence determinants LigA and LigB. Importantly, the PerRA, PerRB and PerRA/B DMC regulons contain multiple genes related to environmental sensing and/or transcriptional regulation. Our data suggest that PerRA and PerRB are part of a complex regulatory network that promotes host adaptation by *L. interrogans* within mammals.

## Introduction

Leptospirosis is a neglected tropical disease with a worldwide distribution [1, 2]. Globally, ~1 million cases and ~60,000 deaths are estimated each year [3]. Leptospirosis is now well recognized as a significant public health problem in developing countries and tropical regions [4–6]. In poor, urban communities in underdeveloped countries, major outbreaks of leptospirosis often are associated with seasonal flooding [4]. Leptospirosis also is of considerable veterinary importance; leptospirosis in cattle and other ruminants can lead to reduced reproductive fitness and diminished milk production, with substantial economic consequences [7, 8].

Leptospirosis is caused by infection with pathogenic spirochetes belonging to the genus *Leptospira* [9]. The majority of severe cases of human leptospirosis are associated with *L. interrogans* [9]. Infection begins when a naïve reservoir (or incidental) host comes into contact with urine from an infected host, most often *via* contaminated water or soil [10]. Leptospires gain entry to the host through bruises or abrasions in the skin and/or mucous membranes. Following inoculation, leptospires transition from a saprophytic (free-living) to a parasitic lifestyle by a complex and poorly understood process referred to as ‘host adaptation’. Once in the bloodstream, leptospires rapidly disseminate to distal tissues but, in reservoir host, are cleared within several days from all sites except the kidney, where they set up long-term residence in the proximal tubules [11–13]. Infected reservoir hosts shed large numbers of leptospires (up to 10^7^/mL) in their urine for weeks to months [9, 12, 14, 15]. While infection in reservoir hosts is generally asymptomatic, incidental hosts, including humans, may develop clinical symptoms ranging from mild flu-like illness to fulminant disease (*e.g.*, Weil’s disease and pulmonary hemorrhage syndrome). Even with treatment, mortality for severe leptospirosis ranges between 10-70% [4]. The factors driving disease severity in humans are poorly understood but are thought to include the bacterial serovar and strain, inoculum size, and the host’s innate and adaptive immune responses [6, 16–18].

The ability of pathogenic *Leptospira* spp. to sense and respond to environmental signals encountered within mammals is generally believed to be critical to sustain the bacterium within its zoonotic lifestyle. The majority of studies investigating gene regulation by *L. interrogans* have done so by manipulating *in vitro* growth conditions [19–27]. However, numerous studies using another enzootic spirochetal pathogen, *Borrelia burgdorferi*, have shown that cultivation *in vitro* under “mammalian host-like” conditions (*i.e.*, increased temperature, increased pH, high osmolality) does not replicate the full range of environmental signals and physiological cues that spirochetes respond to *in vivo* [28–40]. Thus, to gain better insight into the transcriptomic and antigenic changes that *L. interrogans* undergoes within mammals, we developed an *in vivo* model in which leptospires are cultivated within dialysis membrane chambers (DMCs) implanted into the peritoneal cavities of rats, a natural reservoir host [41, 42]. Leptospires within DMCs (6-8 kDa MWCO) are exposed to host-derived nutrients and environmental signals but are protected from the host’s cellular and humoral immune responses. Importantly, the DMC model provides sufficient numbers of host-adapted organisms (~10^8^ per ml) for genome-wide transcriptomics [41] and proteomics [43]. Using this model, we identified 166 genes (110 upregulated and 56 downregulated) differentially-expressed by *L. interrogans* serovar (sv.) Copenhageni strain Fiocruz L1-130 in response to host-specific signals [41]. Almost all of the genes upregulated by the Fiocruz L1-130 strain within DMCs were unique to pathogenic leptospires (*i.e*., not found in the genomes of saprophytic *Leptospira* species).

Not surprisingly, many of the genes upregulated by *L. interrogans* in DMCs encode functions related to environmental signaling and gene regulation [41], including *LIMLP10155* (*LIC12034*), which encodes a member of the <u>F</u>erric <u>U</u>ptake <u>R</u>egulator (FUR) superfamily [44]. The namesake of this highly diverse superfamily, Fur, functions as a global regulator of iron homeostasis in Gram-negative and -positive bacteria, controlling both the induction of iron uptake systems under iron limitation and the expression of iron storage proteins and iron-utilizing enzymes under iron sufficiency [45]. The FUR superfamily is diverse and includes regulatory sensors for zinc (Zur), manganese (Mur) and nickel (Nur) [46–52]. Operating under a divergent regulatory scheme, <u>I</u>ron <u>r</u>esponse regulators (Irrs) sense Fe-heme and repress heme biosynthetic genes under iron-limiting conditions [53]. Unlike most FURs, PerRs, are not involved in metal homeostasis *per se* but instead sense intracellular peroxide and regulate genes involved in detoxification of ROS in a metal-dependent manner [45, 54–58]. While Fur and Zur regulators are widely distributed across both Gram-positive and Gram-negative bacteria, other FUR family regulators have more limited distribution. PerRs are found mainly in Gram-positive bacteria, and Irrs are limited to *α*-proteobacteria [53]. So far, Mur and Nur have been characterized in *α*-proteobacteria and actinomycetes, respectively, but their distribution within other taxonomic groups is still unclear. Typically, FURs act as repressors; inactivation of the regulator leads to ‘constitutive de-repression’ of target genes in the mutant. Although the mechanisms are not well understood, examples of FURs, including PerRs, acting as activators are well documented [59–64]. Beyond balancing metal homeostasis and toxicity, FURs also modulate intermediary metabolism, host colonization and virulence [52, 65–70].

In diverse bacteria, iron serves as an essential co-factor for many cellular processes, including energy generation *via* electron transport, intermediary metabolism and DNA biogenesis [68–72]. For many pathogens, the shift from a high- to low-iron environment is a key environmental signal for induction of expression of virulence genes [69]. Unlike other spirochetes,

such as *B. burgdorferi* and *Treponema pallidum,* which require iron in trace amounts, if at all [73–77], *Leptospira* spp. require this metal for growth *in vitro* and, presumably, in the host [78]. Consequently, leptospires have evolved elaborate mechanisms for iron sensing, scavenging and utilization [79–81]. At the same time, leptospires must balance their physiological need for transition metals with the potential damage caused by highly toxic hydroxyl free radicals generated by Fenton chemistry from H_2_O_2_ in the presence of ferrous ions [82–84]. Many bacteria, including *Leptospira* spp., encode systems to ameliorate the toxicity of H_2_O_2_ and repair damage due to oxidative stress [85, 86]. Expression of gene products involved in oxidative stress responses typically are controlled by one of two master regulators – OxyR and PerR. While OxyR acts as a transcriptional activator for gene products (*i.e.*, catalase and superoxide dismutase) related to detoxification of reactive oxygen species (ROS), PerR acts as a repressor and is released from DNA following exposure to peroxide [48, 54, 87, 88]. Both master regulators respond to similar amounts of H_2_O_2_ [89, 90]; thus, it is unclear why some bacteria have evolved to use PerR while others use OxyR.

A genome-wide survey of *L. interrogans* identified four putative FUR family regulators (*LIMLP04825*, *LIMLP05620*/*perRB*, *LIMLP10155*/*perRA* and *LIMLP18690*). Prior studies by Lo *et al*. [91] and Zavala-Alvarado *et al*. [92], suggest that PerRA functions as a metal-dependent peroxide stress regulator. Consistent with its repressor function in other bacteria, a *L. interrogans perRA* transposon mutant expresses increased levels of catalase, AhpC and cytochrome c-peroxidase and enhanced survival following exposure to peroxide *in vitro* [91–93]. Recently, Zavala-Alvarado *et al*. [93] demonstrated that expression of *perRB* also was increased by H_2_O_2_ *in vitro*. Inactivation of *perRB* increased survival to superoxide but not H_2_O_2_ [93]. These data suggest that PerRA and PerRB likely are functionally distinct. Consistent with this notion, Zavala-

Alvarado *et al*. [93, 94] saw little overlap between the PerRA and PerRB regulons by RNA-Seq analysis of *in vitro*-cultivated organisms. Interestingly, while *perRA* and *perRB* single mutants are virulent in hamsters [91, 94], a *perRA/B* double mutant was avirulent [93]. Collectively, these data argue that *L. interrogans* requires at least one functional PerR-like regulator for infection in mammals.

To investigate the molecular basis for the phenotypic differences between PerRA and PerRB single and double mutants and identify putative virulence-related genes dysregulated by the loss of either/both regulators *in vivo*, we performed comparative RNA-Seq on all three mutant strains cultivated within DMCs. Similar to RNA-Seq data for *in vitro*-cultivated organisms [93, 94], we saw very little overlap between the PerRA and PerRB regulons within mammals. Interestingly, the PerRB DMC regulon was substantially larger than its *in vitro* counterpart [93]. Importantly, by RNA-Seq, we identified 90 genes that are dysregulated only in the double mutant cultivated in DMCs. Of particular note, the “double-only” regulon includes at least four virulence-associated genes, *ligA* and *ligB*, encoding Leptospiral Immunoglobulin-like proteins LigA and LigB, and *lvrAB*, encoding tandem sensory histidine kinases (HKs). Decreased expression of LigA and LigB in the *perRA/B* double mutant was not due to loss of LvrAB signaling. The *perRA/B* double mutant DMC regulon also contains 15 additional genes related to environmental sensing and/or gene regulation, including nine putative hybrid HKs and six putative DNA binding proteins; all but two of the 15 were dysregulated only *in vivo*. Taken together, our data suggest that PerRA and PerRB are part of a complex signaling network that uses mammalian host-specific signals to coordinate the expression of genes required by *L. interrogans* for adaptation to reservoir and incidental (*i.e.*, human) hosts.

## Results

### Pathogenic and saprophytic *Leptospira* spp. encode different FUR-like metalloregulator repertoires

To gain insight into the functions of the four FUR-like regulators encoded by *L. interrogans*, we performed a phylogenetic comparison of these proteins against well characterized representative FUR family metalloregulators from Gram-negative and -positive bacteria (Fig 1A). PerRA and PerRB clustered most closely with PerRs and iron-response regulators (Irrs), while LIMLP18590 and LIMLP04825 clustered with Zur/Nur and Fur/Mur regulators, respectively.

**Fig 1.**
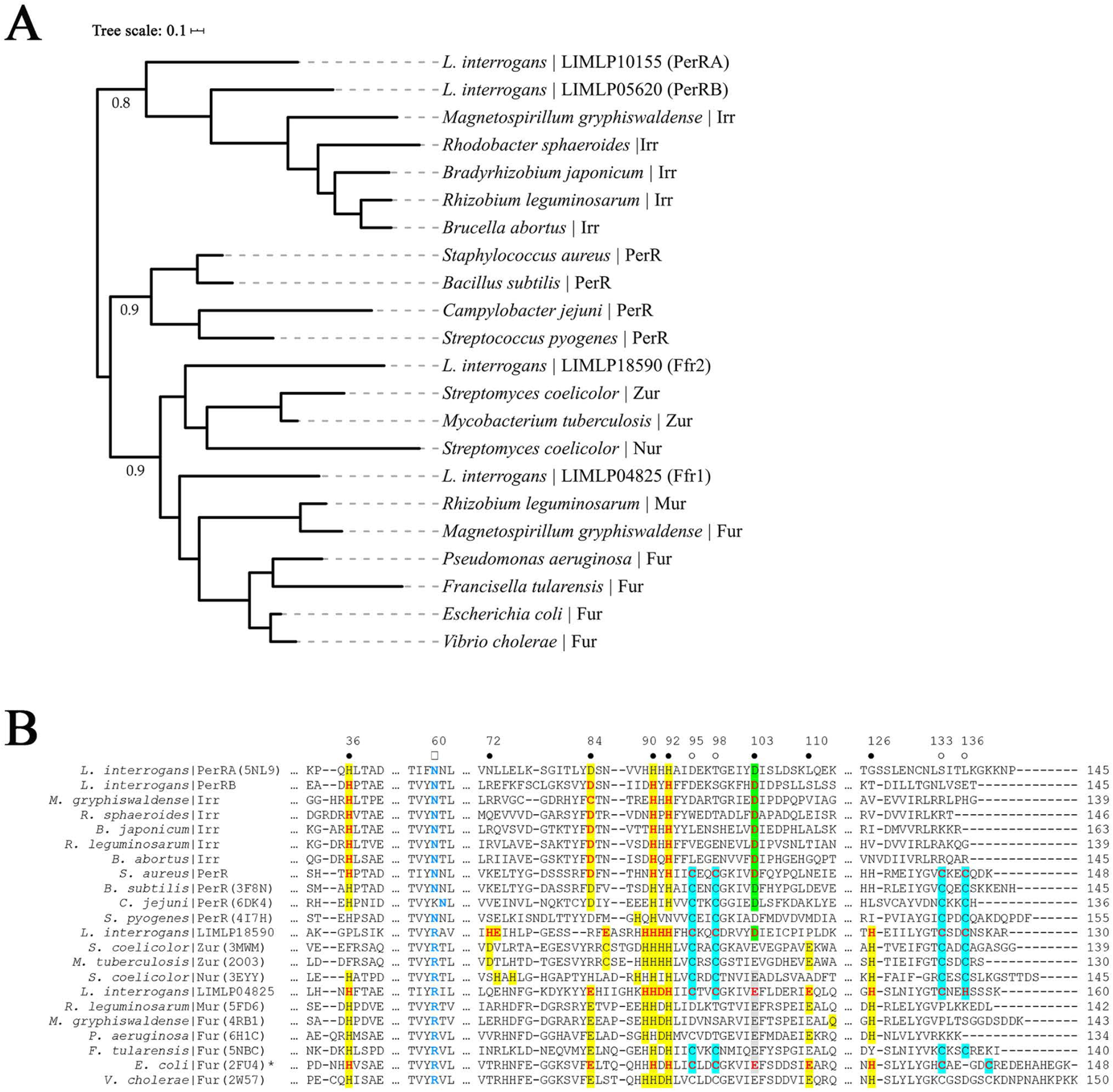
Comparative sequence analysis of Ferric uptake regulator (FUR) domain-containing proteins from *L. interrogans* and other bacteria. **A.** Phylogenetic analysis of *L. interrogans* FUR-like regulators LIMLP10155 (PDB:5NL9, PerRA), LIMLP05620 (PerRB), LIMLP04825 (Ffr1) and LIMLP18590 (Ffr2) with well-characterized FUR superfamily members from diverse bacteria. Phylogenetic analyses were performed as described in Materials and Methods. A midpoint rooted tree was generated using iTOL [201]. Fur family regulators represented in the tree: *Bacillus subtilis* PerR (Uniprot: P71086, PDB: 3F8N); *Bradyrhizobium japonicum* Irr (Uniprot: A0A0A3XTB2); *Campylobacter jejuni* PerR (Uniprot: Q0PBI7, PDB: 6DK4); *Escherichia coli* Fur (Uniprot: P0A9A9, PDB: 2FU4); *Francisella tularensis* Fur (Uniprot: Q5NIN6, PDB: 5NBC); *Magnetospirillum gryphiswaldense* Irr (Uniprot: V6F4I4) and Fur (Uniprot: V6F4Q0, PDB: 4RB1); *Brucella abortus* Irr (Uniprot: Q2YQQ7); *Mycobacterium tuberculosis* Zur (Uniprot: P9WN85, PDB: 2O03); *Pseudomonas aeruginosa* Fur (Uniprot: Q03456, PDB: 6H1C); *Rhizobium leguminosarum* Irr (Uniprot: Q8KLU1) and Mur (Uniprot: O07315, PDB: 5FD6); *Rhodobacter sphaeroides* Irr (Uniprot: Q3IXE0); *Staphylococcus aureus* PerR (Uniprot: Q2G282); *Streptococcus pyogenes* PerR (Uniprot: A0A0H2UT39, PDB: 4I7H); *Streptomyces coelicolor* Zur (Uniprot: Q9L2H5, PDB: 3MWM) and Nur (Uniprot: Q9K4F8, PDB: 3EYY); and *Vibrio cholerae* Fur (Uniprot: P0C6C8, PDB: 2W57). B. Multiple sequence alignment of FUR-like regulators in A. Residues confirmed to be involved in regulatory metal coordination (●) are highlighted in yellow, green or gray; position 103 is used to discriminate between PerR/Irrs (Asp, green) and Fur/Zur/Mur/Nur regulators (Glu, gray). CxxC-motif residues (○) confirmed to be involved in structural metal coordination are highlighted in cyan. Residues in red are predicted but not confirmed by X-ray crystallography to be involved in regulatory or structural metal coordination. Asparagine (N) or arginine (R) residues (1) in blue, located in DNA binding helix H4, can be used to distinguish between PerR and Fur, respectively [97]. *, the PDB structure for *E. coli* Fur includes only the DNA binding domain. Numbers on the top correspond to residues positions in *L. interrogans* PerRA.

We next surveyed the amino acid sequences of the leptospiral FUR-like proteins for conserved regulatory and structural metal binding sites (MBS), which promote DNA binding and folding/dimerization, respectively, in other Fur family regulators [49, 50, 95]. As noted recently by Zavala-Alvarado *et al*. [93], PerRA and PerRB contain two PerR canonical amino acid residues (Asn60 and Asn68 in PerRA and PerRB, respectively) involved in peroxide sensitivity and DNA recognition (Asp103 and Asp112 in PerRA and PerRB, respectively) [96, 97]. Based on these features and increased expression of *perRB* upon exposure of *L. interrogans* to peroxide, LIMLP05620 was named *perRB* [93]. Interestingly, as shown in Fig 1B, the aspartate of the PerR regulatory MBSs and the asparagine in the PerR DNA-binding helices (DBH) also are conserved in Irr proteins. As noted previously by Kebouchi *et al.* [92] and Zavala-Alvarado *et al.* [93, 94], both PerRA and PerRB lack the C-terminal conserved CxxC motif(s) used for structural metal-dependent dimerization by many, but not all, FUR family regulators; this cysteinate motif also is absent in Irrs. Overall, the PerRA and PerRB DBHs are not highly conserved, raising the possibility that they recognize different upstream sequences. LIMLP04825, on the other hand, contains features conserved across Fur, Mur, Zur and Nur regulators, including a glutamic acid at position 103, one or possibly two CxxC motifs (residues 95-98 and 133-136), and an arginine (Arg60) within its DBH (Fig 1B). Interestingly, LIMLP18590 contains features of both PerR (Asp at position 103) and Fur/Mur/Zur/Nur (Arg residue within its putative DBH). The regulatory metal binding site(s) for LIMLP18590 most closely resembles that of a Zur (Fig 1B), which includes two putative tetra-coordinated zinc binding sites rather than the single penta-coordinated site used by PerR. However, without additional data regarding the peroxide responsiveness and/or regulatory metal-binding properties of LIMLP04825 or LIMLP18590, it is not possible to discern their function(s). For this reason, we propose designating them as <u>F</u>ur <u>f</u>amily <u>r</u>egulators 1 (Ffr1) and 2 (Ffr2), respectively.

We next assessed the conservation of FUR family regulators across pathogenic (P1 and P2) and saprophytic (S1 and S2) leptospiral subclades [98]. Orthologs for PerRA and Ffr1 were identified in all highly pathogenic (P1), some intermediate (P2) and all saprophytic strains (S1 and S2), whereas orthologs for PerRB and Ffr2 were found exclusively in pathogenic strains (Figs 2A-B and S1). Our analyses also identified two additional FUR family regulators, both of which were found only in saprophytic leptospires (Figs 2A-B and S1). The first, designated PerRC, contains features of a canonical PerR (two CxxC motifs, an aspartic acid residue within its regulatory MBS and an asparagine within its putative DBH). The second saprophyte-specific FUR family regulator, designated Ffr3, resembles a Fur/Mur/Nur-like regulator (two CxxC motifs, a glutamic acid residue within its regulatory MBS and an arginine within its putative DBH).

**Fig 2.**
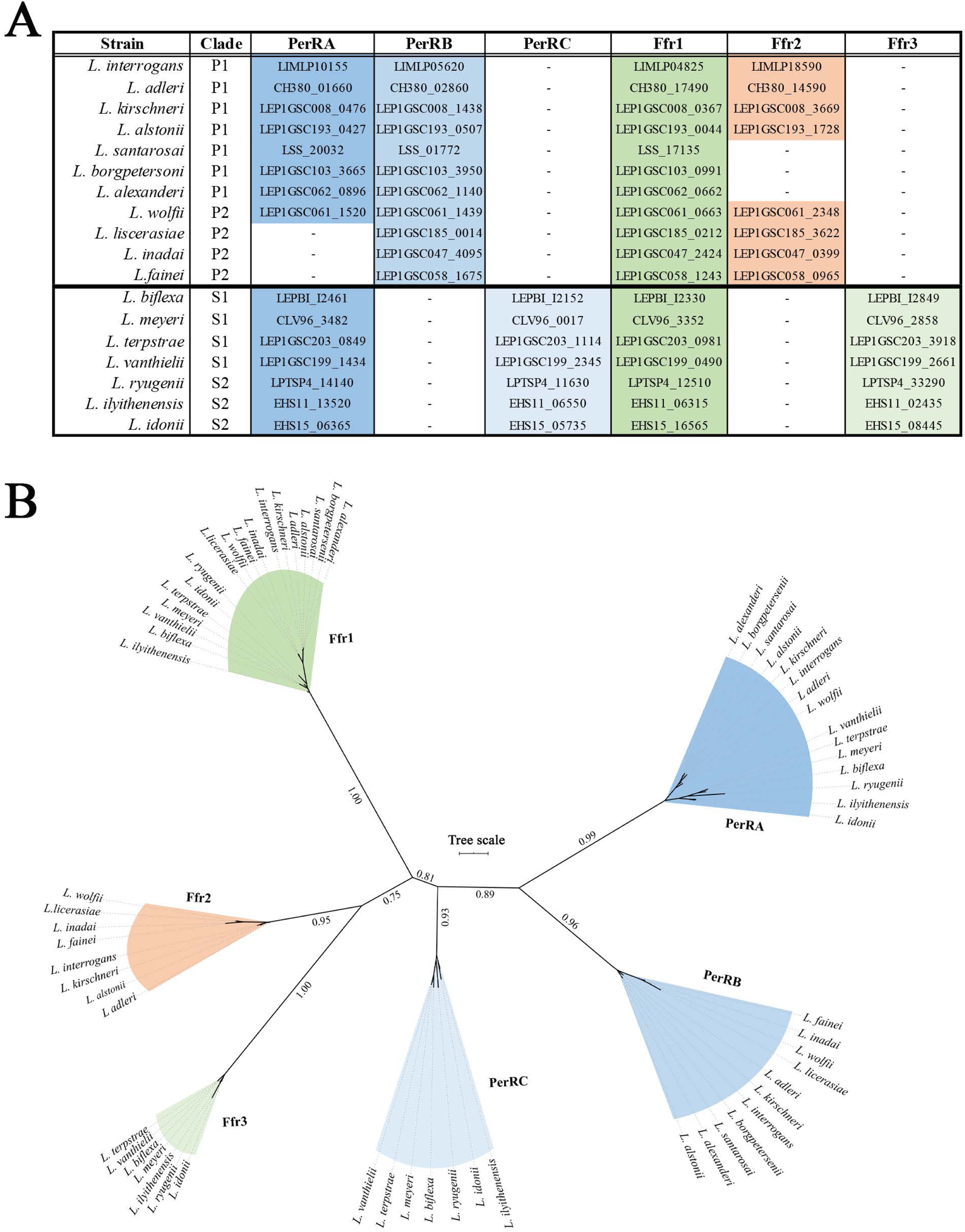
Distribution of Ferric-uptake regulator (FUR) domain-containing proteins across pathogenic and saprophytic *Leptospira* spp. **A.** FUR domain-containing proteins in representative *Leptospira* spp. from pathogenic (P1 and P2) and saprophytic (S1 and S2) subclades. Genomic locus tags for each FUR family protein in *Leptospira* spp. are indicated. **B.** Phylogenetic analysis of *Leptospira* spp. FUR family proteins shown in **A**. Unrooted tree was generated using iTOL [201].

### *L. interrogans* FUR family regulators are expressed at higher or comparable levels in DMCs compared to *in vitro*

Previously, we reported that expression of *perRA* in *L. interrogans* sv. Copenhageni strain Fiocruz L1-130 was induced 3.83-fold in response to mammalian host signals compared to *in vitro* [41]. Using qRT-PCR, we compared transcript levels for all four FUR-like regulators in *L. interrogans* sv. Manilae strain L495 [99, 100] grown *in vitro* (EMJH at 30°C) and following cultivation within DMCs. As shown in Fig 3, *perRA* (7.59-fold), *ffr1* (3.20-fold), and *ffr2* (5.70-fold) were upregulated significantly (*p*<0.05) *in vivo*. *perRB* was upregulated 1.63-fold in DMCs compared to *in vitro*, but the difference was not statistically significant (Fig 3).

**Fig 3.**
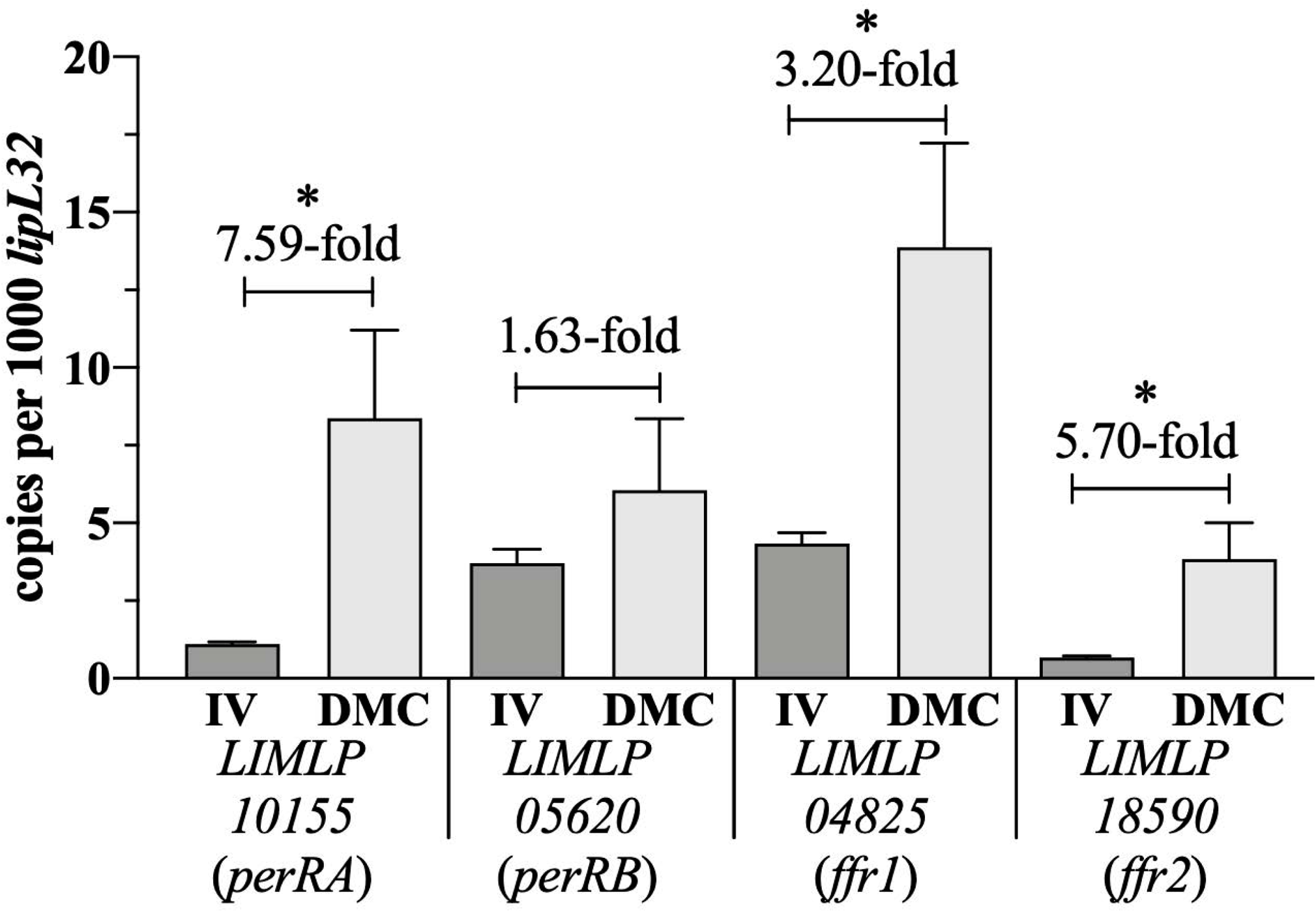
*L. interrogans* express increased transcript levels for three FUR family regulators in response to mammalian host signals compared to *in vitro*. Transcripts for *LIMLP10155* (*perRA*), *LIMLP05620* (*perRB*), *LIMLP04825* (*ffr1*) and *LIMLP18590* (*ffr2*) and were accessed by qRT-PCR using cDNAs from wild-type *L. interrogans* sv. Manilae strain L495 cultivated *in vitro* in EMJH at 30°C (IV) or within rat peritoneal dialysis membrane chambers (DMC). Transcript copy numbers for each gene of interest were normalized per 1000 copies of *lipL32*. Bars show the average of four biological replicates for each condition, assayed in quadruplicate. *p*-values were determined using a two-tailed *t*-test; *, *p* < 0.05.

### Inactivation of both *perRA* and *perRB* in *L. interrogans* results in loss of virulence in mice

Previously, Murray *et al*. [99] and Zavala-Alvarado *et al*. [94] independently reported that a *L. interrogans* Manilae *perRA* Tn mutant is virulent in hamsters. More recently, Zavala-Alvarado *et al.* [93] established that *L. interrogans* lacking PerRB also retain virulence in hamsters. Zavala-Alvarado and colleagues also generated a double mutant by insertional inactivation of *perRA* in the *perRB* Tn mutant; the resulting double mutant (*perRA/B*) was avirulent in hamsters [93]. Golden Syrian hamsters are exquisitely sensitive to *L. interrogans* and develop acute, fulminant, disseminated disease at doses as low as 10^1^ [17, 99, 101, 102]. Mice, on the other hand, are a natural reservoir for *L. interrogans* and relatively resistant to infection; at sublethal doses, susceptible mouse strains develop a self-resolving hematogenous dissemination phase (~1 week) followed by chronic, asymptomatic renal colonization marked by shedding large numbers of leptospires in urine [17, 103–106].

Given the differences in leptospiral disease progression and severity between hamsters and mice, we asked whether PerRA, PerRB, or both are required to establish infection and persistence within a reservoir host model. At the outset, we first established that our wild-type serovar Manilae parent (WT) is virulent in C3H/HeJ mice. Female 10-week old mice (n=5 per group) were infected intraperitoneally with 5 *×* 10^6^, 1 *×* 10^6^, 1 *×* 10^5^ and 1 *×* 10^4^ leptospires. Mice were monitored daily for signs of disease (*i.e.*, weight loss). Within 6 days, all mice in the 5 x 10^6^ group and 3 of 5 mice in the 10^6^ group succumbed to infection, while all others survived the entire 42-day experimental time course (Fig 4A). Based on these virulence studies, the LD_50_ for the wild-type (WT) parent was *≅* 7 x 10^5^. Beginning 14 days post-infection (p.i.), surviving mice were monitored weekly for the presence of leptospires in their urine by darkfield microscopy. Urine from all but one (10^4^ group) mouse contained large numbers of leptospires at all three time points (14, 21, and 35 days p.i.) (Fig 4B). At 42 days p.i., kidneys harvested from all surviving mice infected with the WT parent, including the single urine-negative mouse from the 10^4^ group, were culture-positive.

**Fig 4.**
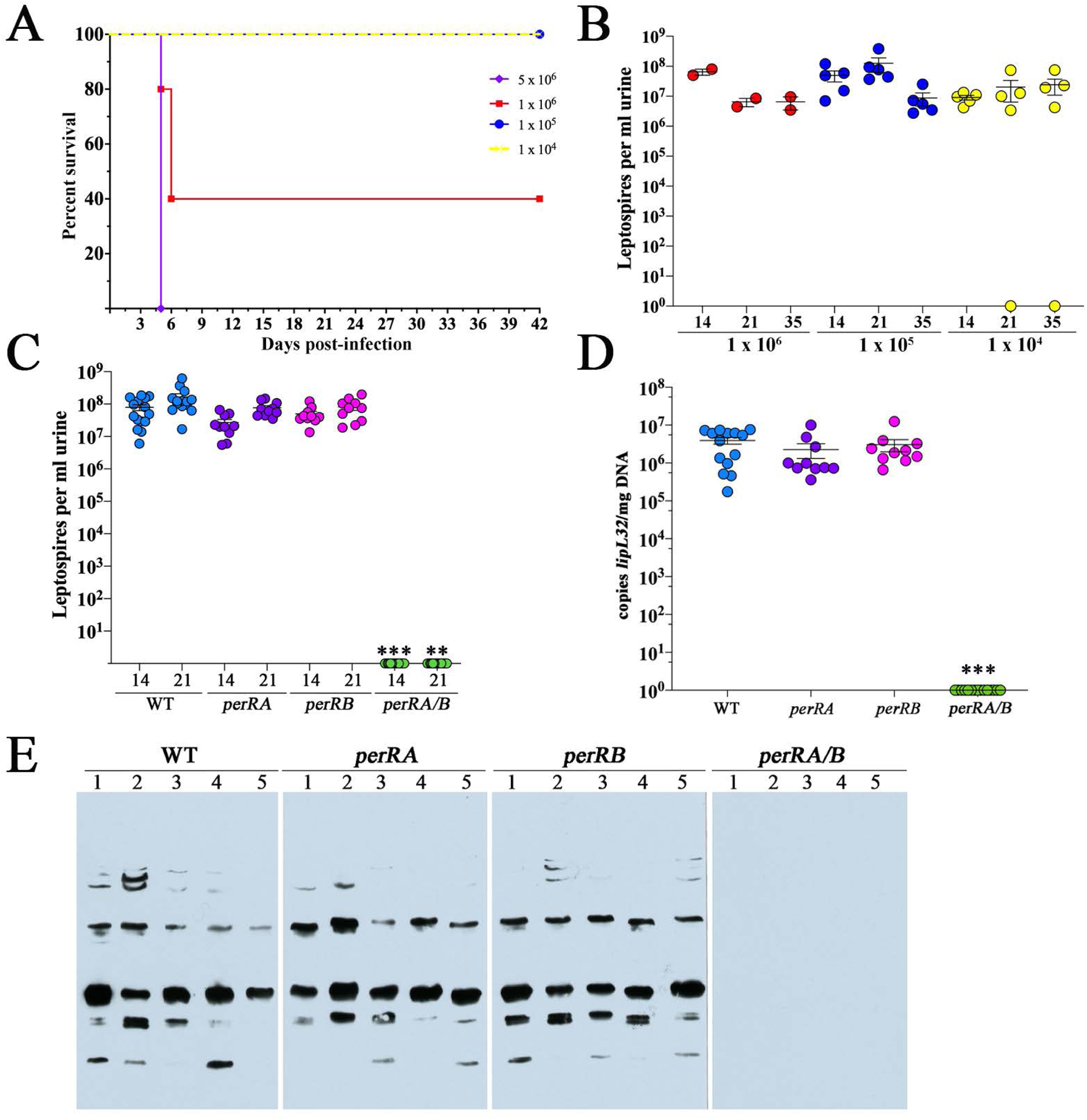
*L. interrogans* requires both PerRA and PerRB for renal colonization of C3H/HeJ mice. **A.** Female 10-week-old C3H/HeJ mice (5 per group) were inoculated intraperitoneally with the indicated numbers of leptospires and monitored for 42 days. At 42 days, animals were euthanized, and kidneys harvested for culturing in EMJH. **B**. Enumeration of leptospires in urine collected from C3H/HeJ mice shown in A 14-, 21- and 35-days post-infection. Circles represent data for urine from individual mice. Burdens per ml of urine were assessed by darkfield microscopy using a Petroff-Hauser counting chamber. Bars show the average and standard error of the mean. *p*-values were determined using a two-tailed *t*-test. **C.** Enumeration of leptospires in urine collected from C3H/HeJ mice inoculated intraperitoneally with 10^5^ of WT, *perRA*, *perRB* or *perRA/B* strains. Burdens per ml of urine were assess by darkfield microscopy using a Petroff-Hauser counting chamber. Circles represent data for urine from individual mice in three independent experiments (5 mice per group, per strain, per experiment). One mouse infected with the WT strain died between day 14-21 due to circumstances unrelated to infection with *L. interrogans*. **D**. Burdens of leptospires in kidneys harvested from mice in panel C. DNA samples from kidneys harvested 28 days post-inoculation were assessed by qPCR using a Taqman-based assay for *lipL32* (in quadruplicate). Bars in B-D represent the average and standard error of the mean. *p*-values in C and D were determined by comparing burdens in mice infected with wild-type (WT) and mutant strains at the same timepoint using a two-tailed *t*-test. ***, *p* ≤ 0.0001; **, *p* = 0.0079. **E**. Immunoblot analysis of sera collected from mice with WT, *perRA*, *perRB* or *perRA/B* strains, collected 28 days post-infection and tested against whole cell lysates of *L. interrogans* sv. Manilae strain L495 grown in EMJH at 30°C.

Prior to using the *perRA* and *perRB* single and double mutants for murine virulence studies, we first confirmed their genotypes by amplicon sequencing using primers listed in S6 Table and immunoblot and established that loss of one regulator had no obvious effect on expression of the other in the corresponding single mutants (S2 Fig). We next compared infectivity of the WT, *perRA, perRB* or *perRA/B* strains in C3H/HeJ mice using a sublethal intraperitoneal inoculum (1 x 10^5^). All of the mice inoculated with the WT parent and single mutants were infected, shedding comparable numbers of leptospires in their urine at 14- and 21-days p.i. (Fig 4C). In contrast, no leptospires were detected in urine from mice inoculated with the *perRA/B* double mutant. Consistent with data from urine, at day 28 p.i., all *perRA/B-*infected mice were negative for leptospires by both culture and qPCR (Fig 4D). Lastly, in contrast to mice infected with the WT or single mutant strains, all of which generated robust serological responses against *L. interrogans*, none of the mice infected with the double mutant seroconverted (Fig 4E).

### *perRA* and *perRB* single and double mutants grow normally in rat peritoneal cavities

PerR regulators have been linked to a wide range of physiological functions outside of oxidative stress, including metal homeostasis, metabolism and virulence [107–109]. In *Bacillus subtilis*, inactivation of *perR* leads to increased expression of *fur* and iron starvation [82]. To examine whether the avirulent phenotype of the *perRA/B* double mutant could be due to an inability to grow in mammals, we took advantage of our DMC model, whereby leptospires are cultivated for 9-10 days within dialysis membrane chambers implanted in the peritoneal cavity of a rat [41, 110]. Originally developed for *B. burgdorferi*, this model is able to separate genes related to physiological adaption (*i.e.*, nutrient acquisition and metabolism) from those encoding virulence determinants, such as adhesins, motility and immune evasion. However, we saw no significant difference (*p*>0.05) in the mean number of leptospires for the wild-type (1.75 *×* 10^8^/ml) strain versus each mutant (*perRA,* 8.5 *×* 10^7^/ml; *perRB*, 3.57 *×* 10^8^/ml; and *perRA/B,* 2.70 *×* 10^8^/ml) recovered from DMCs 9 days post-implantation (3 biological replicates per strain). These data demonstrate that the virulence-defect observed with the double mutant is not due to a metabolic lesion (*i.e*., metal starvation).

### Defining the PerRA and PerRB regulons *in vivo* by comparative RNA-Seq

Prototypical FUR family regulators, including PerR, modulate transcription by binding to DNA *via* one or more ~19-bp inverted repeats (‘boxes’) located upstream of their target genes [111]. Kebouchi *et al*. [92] previously identified three potential PerR binding sites upstream of *perRA* in *L. interrogans*. However, searches of the Manilae genome using these sequences, as well as canonical Fur and PerR boxes [111, 112], did not identify additional hits [41, 44, 91]. Therefore, to identify genes controlled by PerRA, PerRB, or both, in response to host signals, we performed comparative RNA-Seq using WT, *perRA*, *perRB* and *perRA/B* strains cultivated in DMCs (3 biological replicates per strain); a summary of the raw Illumina read data is presented in S1 Table. Reads were mapped using EDGE-pro [113] and analyzed for differentially-expressed genes using DESeq2 [114]. Genes expressed at *≥*3-fold higher/lower levels in the WT versus mutant with a False-discovery rate (FDR)-adjusted-*p* value (*q*) *≤*0.05 were considered differentially expressed. Complete RNA-Seq datasets for all comparisons are presented in S2-S4 Tables. Raw read files have been deposited in the NCBI Sequence Read Archive (SRA) database (BioProject accession PRJNA659512).

#### Overview of the PerRA DMC regulon

The PerRA DMC regulon contained a total of 81 differentially expressed genes; 43 were expressed at higher levels (*i.e*., upregulated directly or indirectly by PerRA) in the WT parent compared to the *perRA* mutant, while 38 were expressed at lower levels (*i.e*., downregulated/repressed directly or indirectly by PerRA) (S2 Table). Notably, the PerRA DMC regulon is substantially larger than its *in vitro* counterpart (17 genes total but only 14 dysregulated >3-fold), recently reported by Zavala-Alvarado *et al.* [94]. Overlap between the PerRA DMC and *in vitro* regulons consists primarily of seven genes located in a single chromosomal locus (S3 Fig) containing LipL48 (*LIMLP04280*), a putative outer-membrane embedded TonB-dependent receptor (TBDR, *LIMLP04270*) and one of the 2-3 putative TonB/ExbD/ExbB transporters systems (*LIMLP04245-04230*) encoded by *L. interrogans* [44, 91]. TonB-dependent transporters (TBDT) for iron typically are repressed by Fur [115]; thus, it was surprising that this system was upregulated by PerRA both *in vitro* and in DMCs. Interestingly, none of the prototypical oxidative stress-related genes identified by Zavala-Alvarado *et al.* [94] as being under PerRA control *in vitro* were dysregulated in DMCs.

More than half (55%) of genes in the PerRA DMC regulon encode proteins of unknown function (S5A Fig). The remaining genes are distributed over a wide range of functional categories (COGs) related to cellular homeostasis and metabolism. Most notably, the PerRA regulon includes five genes (two upregulated, three downregulated) involved in signaling and/or gene regulation (Figs 5A and S4A and S2 Table). The three upregulated genes (*LIMLP02515*, *LIMLP05780* and *LIMLP01845*) encode putative DNA binding proteins, including a CsoR-like metal sensitive repressor, while the three downregulated genes encode a two-component system (TCS) histidine kinase with four Per-Arnt-Sim (PAS)-type sensor domains (*LIMLP10140*), a putative DNA binding protein (*LIMLP00900*) and a putative serine/threonine kinase with GAF domain (*LIMLP11575*). PAS domains are ubiquitous in bacteria and sense a wide range of ligands, including heme, FAD, fatty acids and divalent metals [116, 117]. GAF domains share a similar fold to PAS domains and often regulate the catalytic activity of cyclic nucleotide phosphodiesterases [118]. Of note, none of these putative regulatory factors were dysregulated ≥3 fold by loss of PerRA *in vitro* [94].

**Fig 5.**
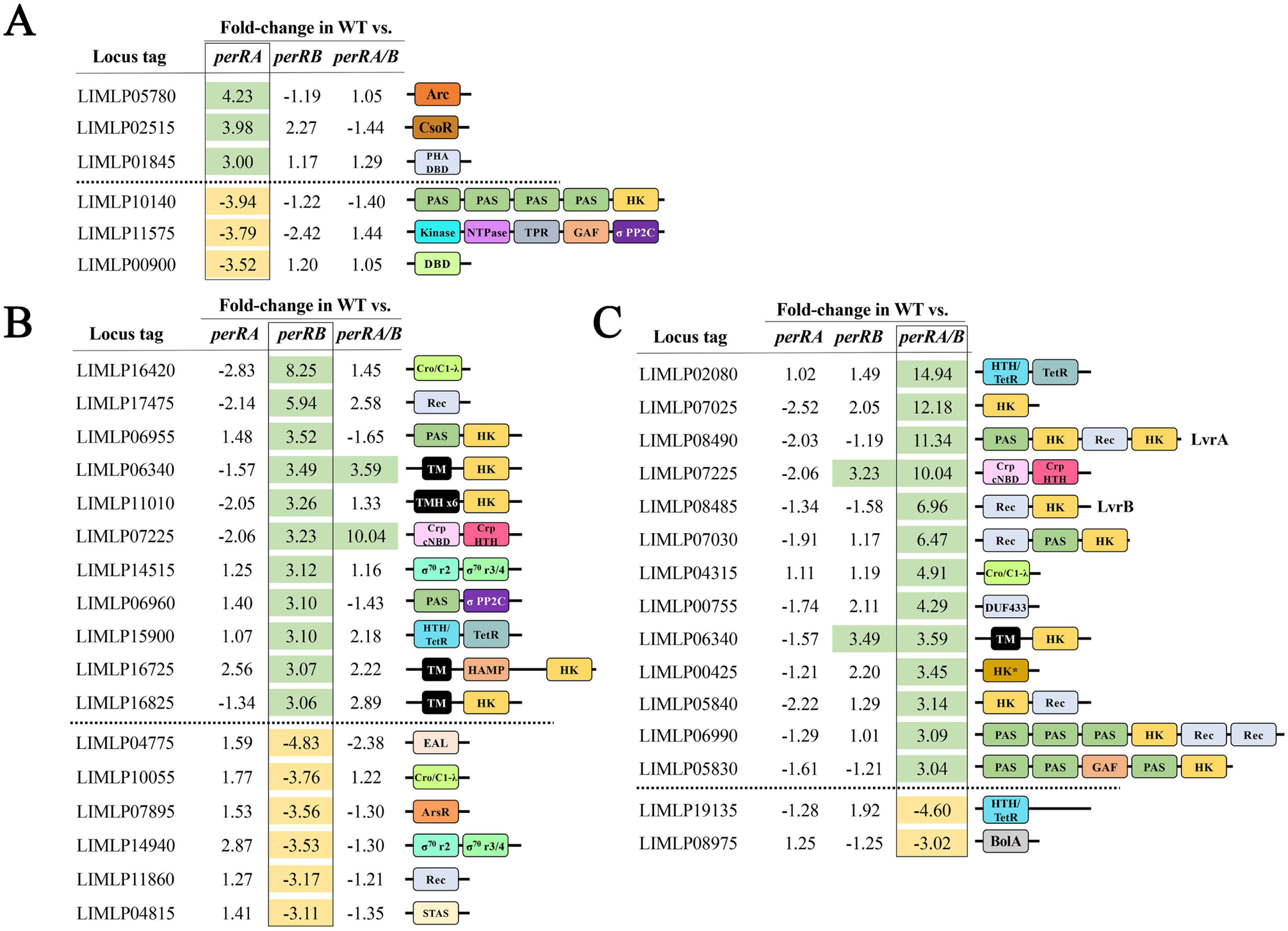
The PerRA and PerRB DMC regulons contain genes related to environmental sensing, signaling and/or transcriptional regulation. Proteins containing conserved domains related to signal transduction systems, DNA binding or other regulatory functions identified as being differentially expressed in the wild-type (WT) vs. *perRA* (**A**), *perRB* (**B**) or *perRA/B* (**C**) RNA-Seq comparisons. Values indicate fold-regulation (up or down) in each comparison. Shading indicates genes differentially expressed ≥3-fold (adjusted-*p* <0.05). Abbreviations for conserved domain names and Interpro (IPR) designations: *σ* PP2C, PPM-type phosphatase domain superfamily (IPR036457); *σ*^70^ r2, RNA polymerase sigma factor region 2 (IPR007627); *σ*^70^ r3/4, RNA polymerase sigma factor, region 3/4-like (IPR013324); Arc, Arc-type ribbon-helix-helix (IPR013321); ArsR, ArsR-type helix-turn-helix DNA-binding domain (IPR001845); BolA, BolA family domain (IPR002634); Cro/C1-*λ*, Cro/C1-type helix-turn-helix domain (IPR001387) and/or Lambda repressor-like, DNA-binding superfamily domain (IPR010982); Crp cNBD, Cyclic nucleotide-binding domain (IPR000595); Crp HTH, Crp-type helix-turn-helix domain (IPR012318); DBD, Putative DNA-binding domain superfamily (IPR009061); DUF433, domain of unknown function DUF433 (IPR007367) and Homeobox-like superfamily domain (IPR009057); EAL, EAL-type phosphodiesterase domain (IPR001633); GAF, GAF-like domain superfamily domain (IPR029016); HAMP, HAMP domain (IPR003660); HK, Histidine kinase (IPR005467, IPR003594) and dimerization/phosphoacceptor (IPR003661) domains; HK*, Histidine kinase (IPR005467, IPR003594) only (no dimerization domain); HTH/TetR, DNA-binding helix-turn-helix/TetR-type domain (IPR001647); Kinase, Protein kinase domain (IPR000719); NTPase, P-loop containing nucleoside triphosphate hydrolase (IPR027417); PAS, PAS domain (IPR000014); PHA DBD, PHA accumulation regulator DNA-binding, N-terminal (IPR012909); REC, Signal transduction response regulator receiver (IPR001789) and/or CheY-like superfamily (IPR011006) domain; STAS, Sulphate Transporter and Anti-Sigma factor antagonist domain (IPR002645); TetR, Tetracyclin repressor-like superfamily C-terminal domain (IPR036271); TM, transmembrane helix; TMx6, six transmembrane helices; and TPR, Tetratricopeptide-like helical domain superfamily (IPR011990).

#### Overview of the PerRB DMC regulon

Inactivation of *perRB* resulted in dysregulation of 200 genes (131 upregulated and 69 downregulated) within DMCs (S3 Table). In contrast, only 30 genes were dysregulated in the *perRB* mutant *in vitro*, with only one affected >3-fold [93]. Remarkably, we saw no overlap between the *in vitro* and DMC PerRB regulons. Overlap between the PerRA and PerRB DMC regulons was limited to genes within the TonB-dependent transporter locus described above (S3 Fig). Notably, none of the TonB-related genes were dysregulated ≥3 fold in the *perRB* mutant *in vitro* (S3 Fig). The implications of these data are two-fold; differences between the *in vitro* and DMC regulons for the *perRB* mutant imply that PerRB is not fully activated under normal growth conditions *in vitro*, while the minimal overlap between the PerRA and PerRB DMC regulons suggests that they recognize different upstream binding sites.

The majority (66%) of genes in the PerRB DMC regulon were upregulated; most of these encode proteins with unknown or poorly characterized functions (S4B Fig and S3 Table). Notably, however, the PerRB DMC regulon includes 17 genes (11 upregulated, 6 downregulated) related to signaling and/or gene regulation (Fig 5B). The 11 upregulated genes include six related to signal transduction, three putative DNA binding proteins (*LIMLP16420*, *LIMLP07225* and *LIMLP15900*), an ECF-type sigma factor (*LIMLP14515*) and a putative serine/threonine phosphatase with a PAS-type sensor domain (*LIMLP06960*) (Fig 5B). The six downregulated signaling genes include two additional putative DNA binding proteins (*LIMLP07895* and *LIML10055*), a second ECF-type sigma factor (*LIMLP14940*), a putative anti-sigma factor antagonist (*LIMLP04815*), and an EAL-type phosphodiesterase (*LIMLP04775*) (Fig 5B). None of these putative regulators were affected *in vitro* by loss of PerRB [93].

### Inactivation of both PerRA and PerRB results in a DMC regulon that differs dramatically from its single mutant counterparts

The PerRA/B DMC regulon contains 106 differentially expressed genes, 74 upregulated and 32 repressed (Tables 1–2 and S4). Surprisingly, we saw limited overlap between the DMC regulons for the double and single mutants (Fig 6A); all of the overlapping genes were located in the TonB-related chromosomal locus dysregulated in the *perRA* and *perRA/B* mutants *in vitro* [93, 94] (S3 Fig). Ninety genes (62 upregulated and 28 repressed) were dysregulated <u>only</u> in the *perRA/B* double mutant (Fig 6A and Tables 1–2 and S4).

**Fig 6.**
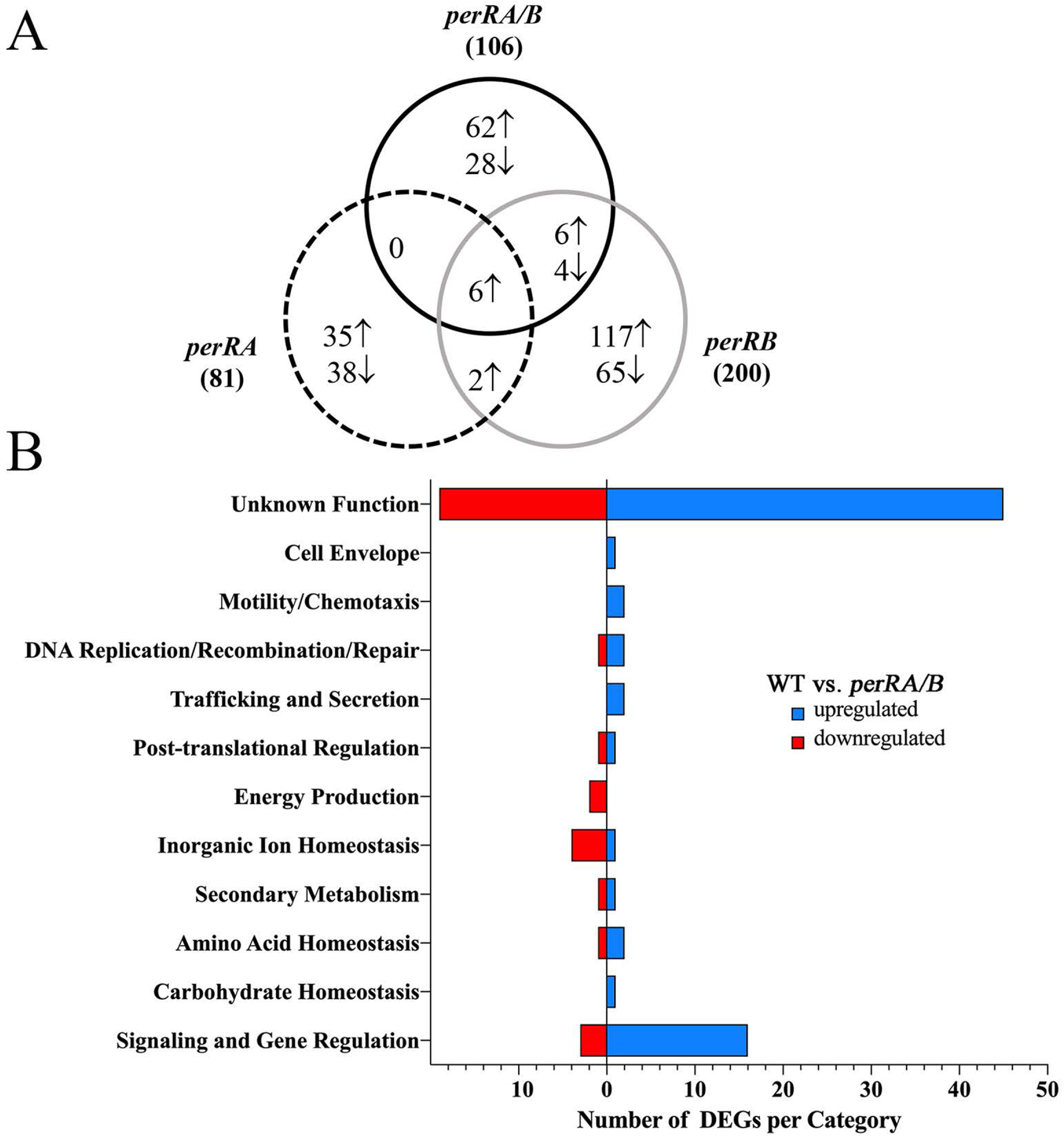
Overview of comparative RNA-Seq analyses of *L. interrogans* wild-type, *perRA*, *perRB* and *perRA/B* mutant strains. **A.** Venn diagram showing overlap of genes differentially expressed *≥*3-fold (*q <u>≤</u>*0.05) in wild-type versus single and double mutant comparisons. *↑* and *↓* symbols denote genes upregulated (*i.e.*, expressed at higher levels in the wild-type vs. mutant) or downregulated (*i.e.*, expressed at lower levels in the wild-type vs. mutant), respectively, by PerRA, PerRB or both (PerRA/B). Complete datasets of all comparisons are presented in S2-S4 Tables. **B.** Cluster of Orthologous Genes (COG) categorization of differentially expressed genes (DEGs) in the wild-type vs. *perRA/B* double mutant RNA-Seq comparison. COG predictions for individual genes are presented in S4 Table. Number of DEGs in each COG are indicated on the *x*-axis.

**Table 1.**
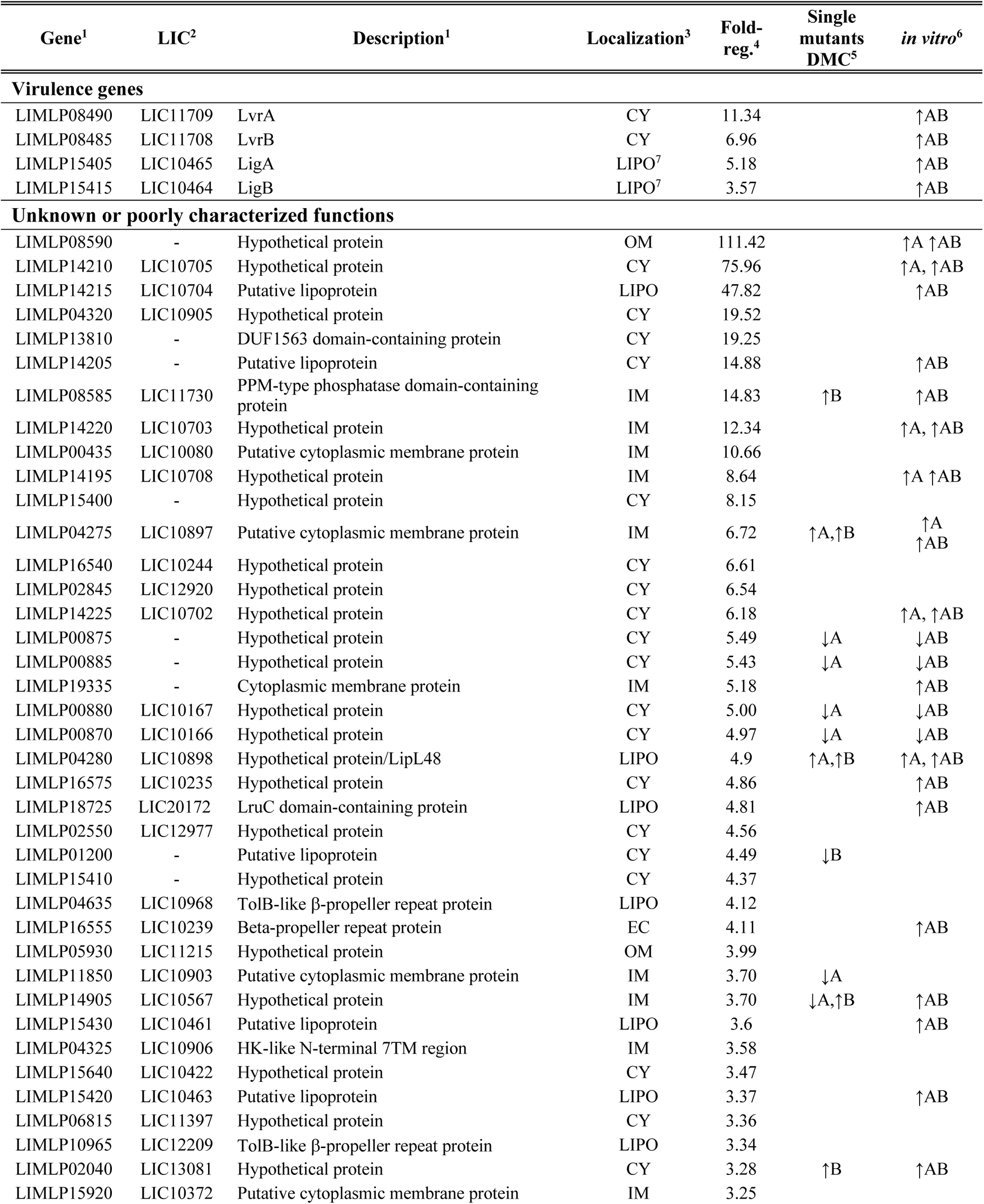
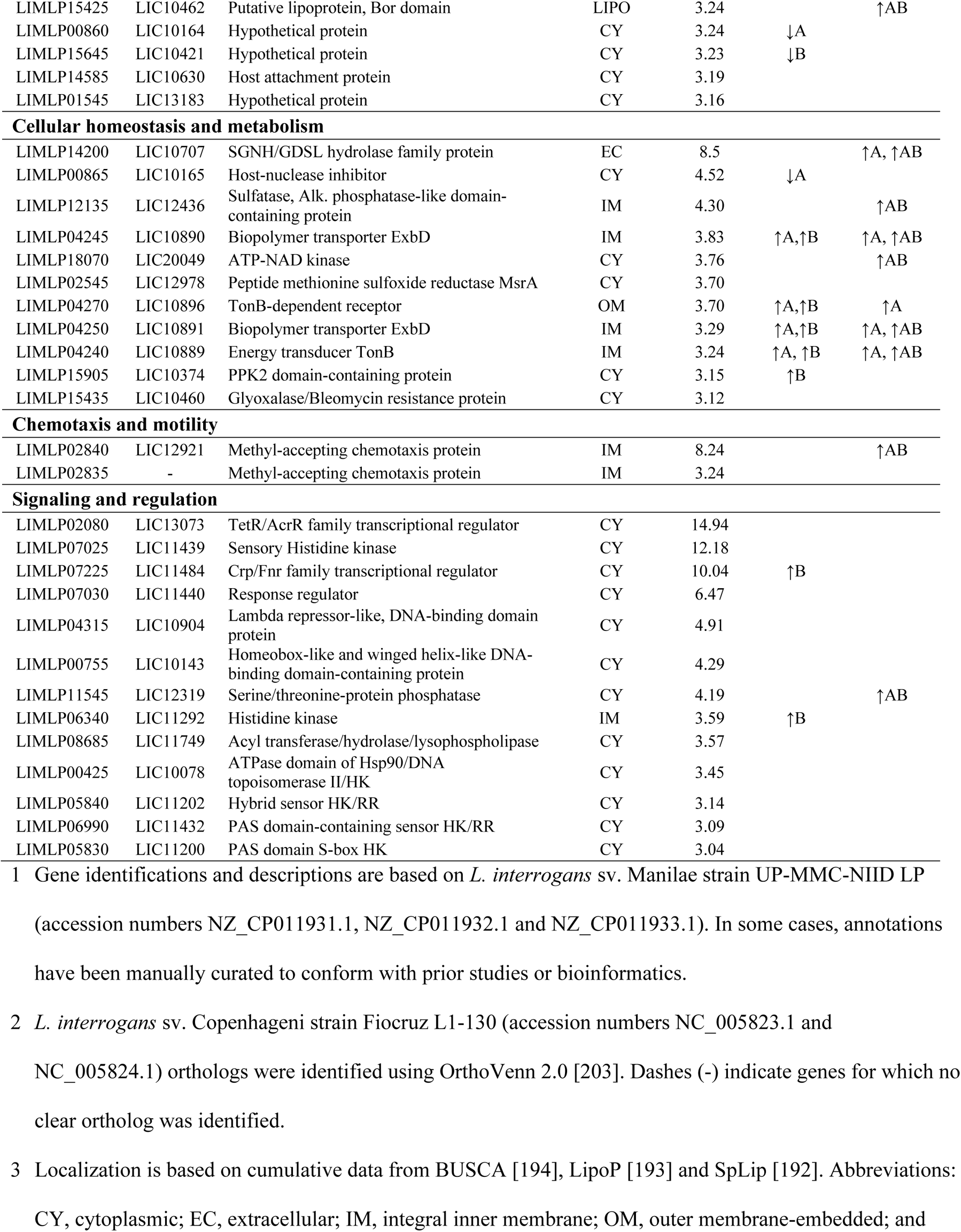
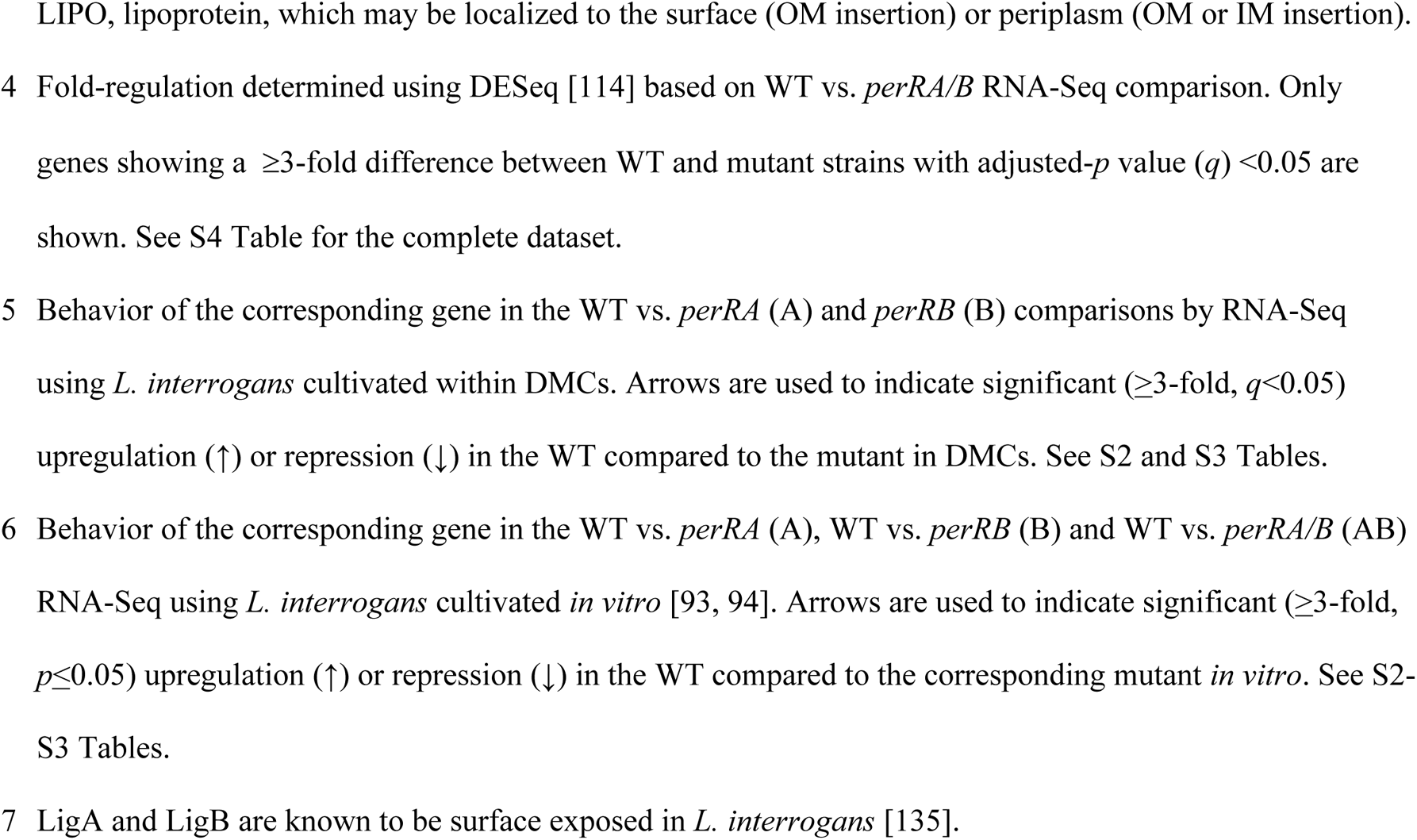
Genes significantly upregulated by PerRA/PerRB in L. interrogans cultivated within DMCs.

**Table 2.**
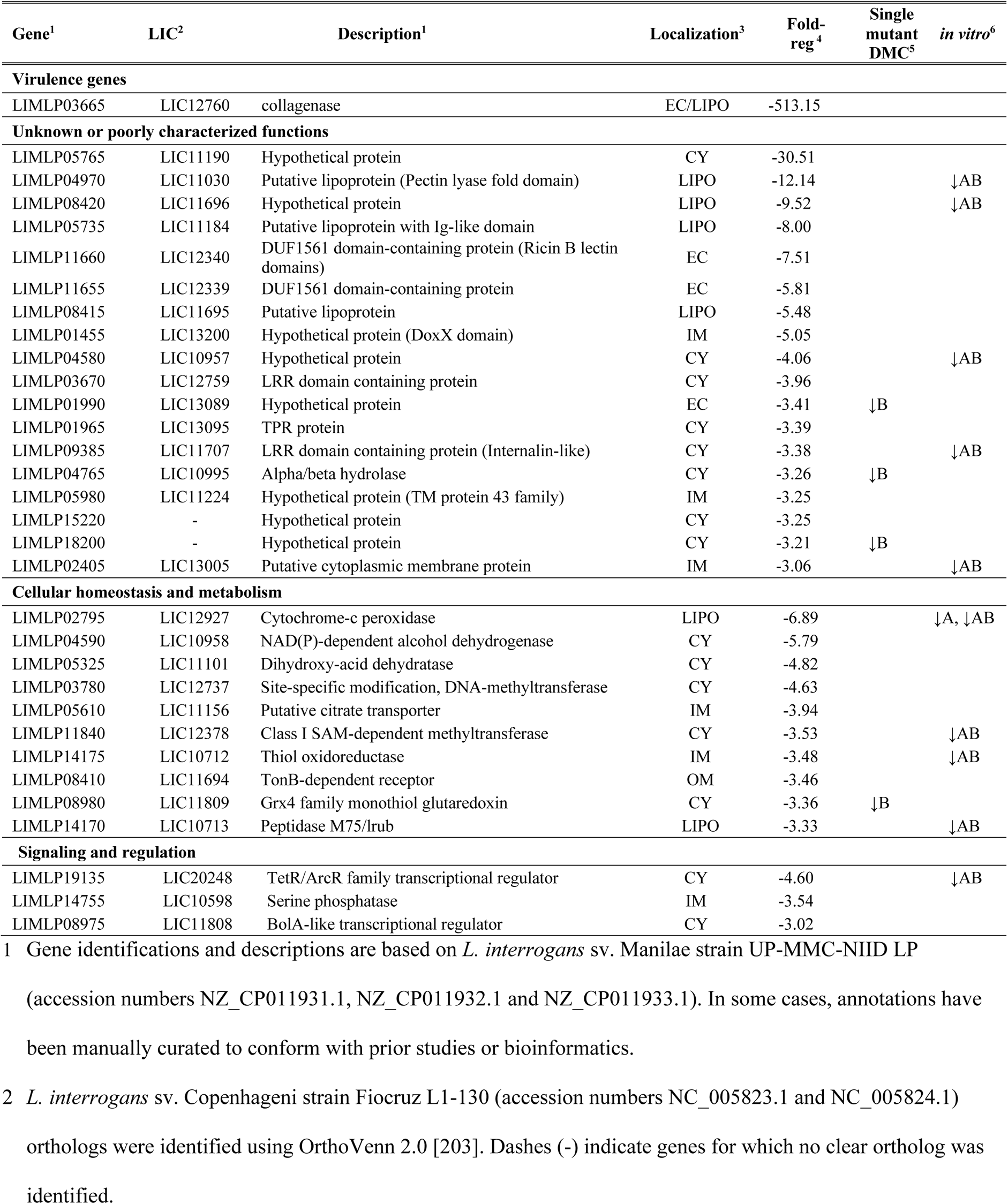
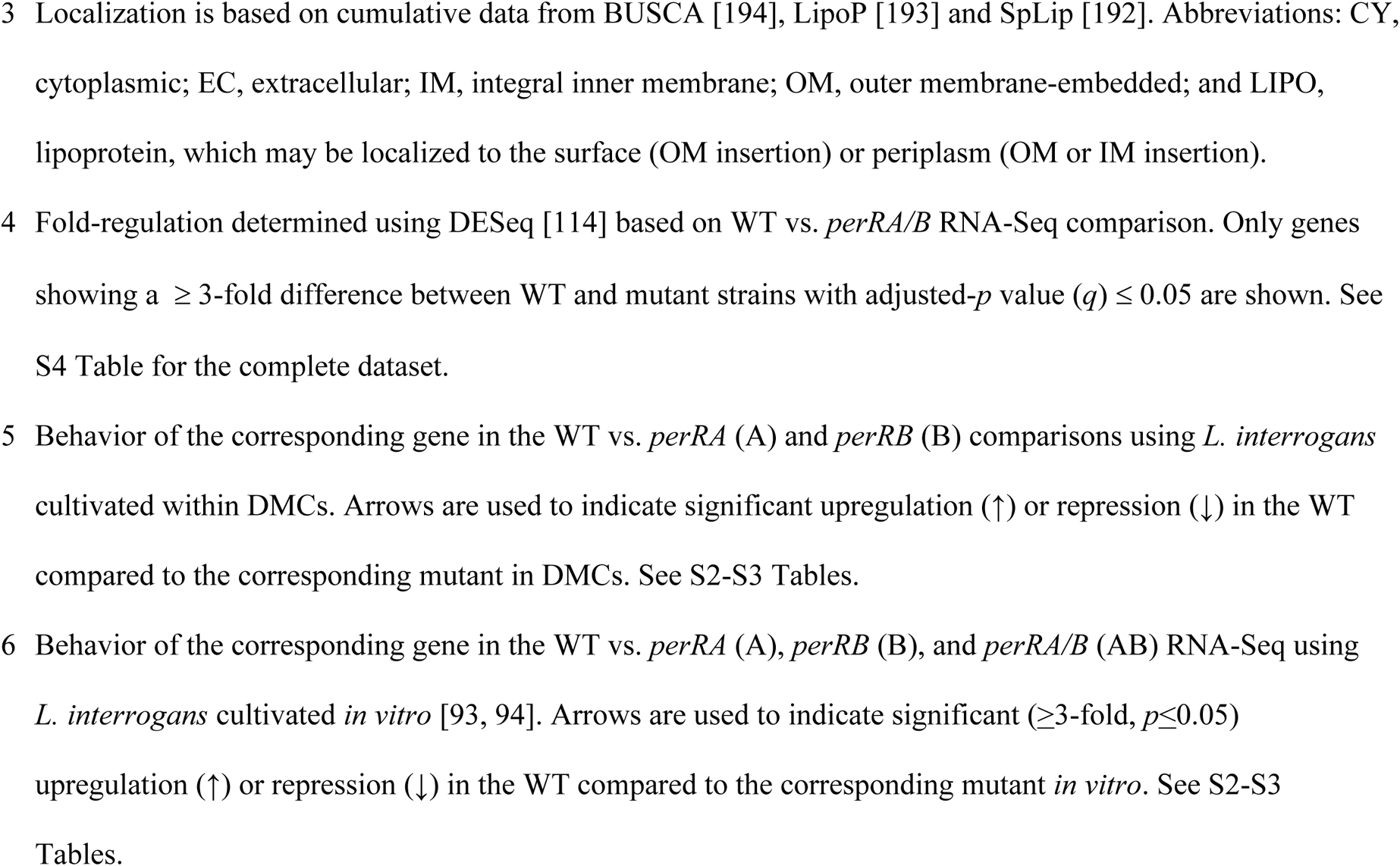
Genes significantly downregulated by PerRA/PerRB in *L. interrogans* cultivated within DMCs.

#### Cellular homeostasis and metabolism

A handful of genes upregulated by PerRA/B in DMCs encode proteins involved in cellular homeostasis and metabolism (Fig 6B and Table 1). *LIMLP14200* and *LIMLP12135* contain domains found in lipases/esterases (IPR0002489) [119] and alkaline phosphatases and sulfatases (IPR000917), respectively. *LIMLP18070* contains an ATP-NAD kinase domain (IPR022504), suggesting a role in maintaining NADP homeostasis and, by extension, NADPH-dependent reductive biosynthetic pathways. *LIMLP02545*, encoding one of the few gene products in the PerRA/B regulon related to oxidative stress, is a putative methionine sulfoxide reductase, which catalyzes the reversible thioredoxin-dependent oxidation-reduction (repair) of Met-SO to Met [120, 121]. Lastly, *LIMLP15435* contains a domain found in glyoxalase/bleomycin resistance proteins; in bacteria, glyoxalases are used to detoxify methylglyoxal, a reduced derivative of pyruvate, as part of the glutathione-dependent glyoxalase system [122].

Eleven genes downregulated by PerRA/B in DMCs are involved in cellular homeostasis and metabolism (Fig 6B, Tables 2 and S4). Only three (*LIMLP02795*, *LIMLP14175* and *LIMLP08980*), encoding a cytochrome c peroxidase, a thiol oxidoreductase and a Grx4 family monothiol glutaredoxin, respectively, are involved in oxidative stress adaptation.

#### Sensing and responding to the mammalian host environment

The PerRA/B regulon includes at least 17 genes related to environmental sensing, signaling and, potentially, host adaptation. Two (*LIMLP02835* and *LIMLP02840*) encode methyl-accepting chemotaxis proteins co-regulated with *LIMLP02845*, encoding a small (62 aa) hypothetical protein of unknown function (Table 1). Nine, including *lvrAB* (discussed below), encode sensory histidine kinases, most of which contain PAS-type sensor domains (Fig 5C). One of the nine (*LIMLP05830*) encodes a regulator that contains both PAS and GAF domains (Fig 5C). The PerRA/B DMC regulon includes six putative DNA binding proteins, four upregulated and two downregulated (Fig 5C). Three upregulated genes belong to the TetR (*LIMLP02080*), Cro/C1-*λ* (*LIMLP04315*) and CRP-like (*LIMLP07225*) repressor families, while the fourth (*LIMLP00755*) encodes a hypothetical protein containing a homeobox winged helix-like domain of unknown function (DUF433). A second TetR-like repressor (*LIMLP19135*) and a BolA-like regulator (*LIMLP08975*) were repressed by PerRA/PerRB in DMCs (Fig 5C). In *E. coli*, BolA has been linked to a range of adaptive responses, including biofilm formation and entry into stationary phase [123]. All but two of the regulatory proteins in the PerRA/B DMC regulon were dysregulated only in the double mutant; *LIMLP06340*, encoding a histidine kinase, and *LIMLP07225*, encoding a CRP-like DNA binding protein, also were upregulated by PerRB alone in DMCs (Fig 5B).

Although the vast majority (70%) of genes upregulated by PerRA/B encode proteins of unknown function (Fig 6B and Table 1), seven contain conserved domains potentially related to mammalian host adaptation and/or virulence. *LIMLP08585* contains a PPM-type phosphatase domain (IPR001932); PPM domains are found in diverse regulatory proteins, including SpoIIE in *B. subtilis* [124]. *LIMLP15425* contains a putative Lambda_Bor-like domain (PF06291), which in *E. coli* has been associated with increased serum survival [125, 126]. *LIMLP14585*, annotated as a host attachment protein, contains a domain of unknown function (IPR019291) found in virulence-associated proteins from the plant pathogens *Agrobacterium tumefaciens* and *Xanthomonas* spp. [127, 128]. *LIMLP02040* contains a SRPBCC-like domain (cd07812), which forms a deep, hydrophobic ligand binding pocket capable of binding diverse ligands [129, 130]. Three hypothetical proteins (*LIMLP04635*, *LIMLP10965* and *LIMLP16555*) upregulated by PerRA/B are predicted to form *β*-propeller structures, which are associated with a wide range of functions, including ligand-binding, enzymatic activity, cell signaling, and protein-protein interactions [131]. Interestingly, Thibeaux et al. [132] previously noted that proteins with *β*-propeller repeats are enriched in highly virulent *Leptospira* spp. Six upregulated genes encode uncharacterized lipoproteins of unknown function (Table 1).

Eighteen (56%) genes downregulated by PerRA/B in DMCs encode proteins of unknown function (Table 2). *LIMP04970* and *LIMLP11660*, both predicted to encode lipoproteins, contain domains (pectin lyase-fold/IPR011050 and Ricin B lectin/IPR000772, respectively) potentially involved in binding to and/or cleavage of host-derived carbohydrates. *LIMLP04765* contains an alpha/beta hydrolase domain shared by a wide range of hydrolytic enzymes. Lastly, *LIMLP01455*, encoding an inner membrane protein, contains a DoxX-like domain; in *Mycobacterium tuberculosis*, DoxX complexes with a thiosulfate sulfurtransferase (SseA) to promote resistance to agents that disrupt thiol homeostasis [133].

#### Known or putative virulence determinants

The upregulated portion of the PerRA/B regulon contains at least four virulence-associated genes (Table 1). Two, *LIMLP15405/ligA* and *LIMLP15415/ligB,* encode the pathogen-specific, multifunctional, <u>L</u>eptospiral <u>I</u>mmuno<u>g</u>lobulin-like repeat proteins LigA and LigB, respectively [134, 135], while *LIMLP08490* and *LIMLP08485* encode the hybrid histidine kinases LvrA and LvrB, respectively [19]. Although tandemly located on the chromosome, *ligA* and *ligB* are not co-transcribed (Fig 7A). They do, however, have identical upstream regions and respond similarly *in vitro* to conditions used to mimic the mammalian host milieu (*e.g.,* high osmolality and increased temperature) [21–24, 136]. Three genes located downstream of *ligB*, all encoding hypothetical proteins, also were upregulated (Fig 7A). Using antisera against the shared N-terminal repeats (Fig 7A), we compared expression of LigA and LigB in WT and mutant strains. As shown in Fig 7B, both LigA and LigB were completely absent in whole cell lysates prepared from the *perRA/B* double mutant cultivated within DMCs (Figs 7B and S5A-B); expression of both Ligs was restored to near wild-type levels by *trans*-complementation with *perRB* alone (Figs 7B and S5A-B). Interestingly, we saw a modest to substantial reduction in LigA/LigB in the *perRA* and *perRB* single mutants. (Figs 7B and S5A-B). Given that the upstream regions for *ligA* and *ligB* are identical, the molecular basis (e.g., transcriptional, post-transcriptional or both) for the difference between Lig levels in the *perA* and *perRB* mutants is unclear.

**Fig 7.**
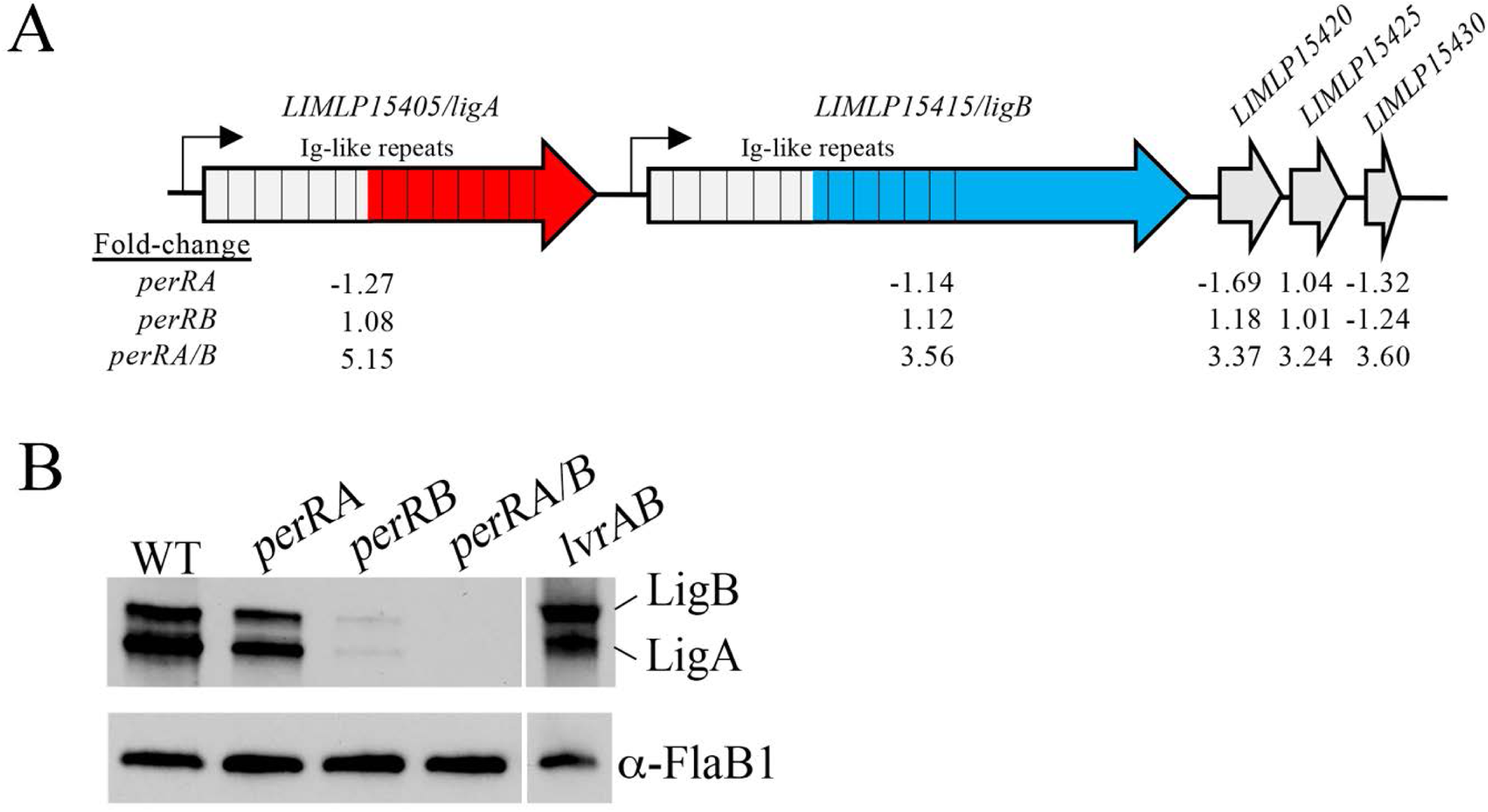
Expression of LigA and LigB *in vivo* is reduced in perRA and perRB single mutants and undetectable in only in the *perRA/B* double mutant. Cartoon depiction of *ligA*, *ligB* and surrounding genes in *L. interrogans* sv. Manilae strain L495. Hatched bars are used to show Immunoglobulin-like repeats. Red and Blue colored regions indicate LigA- and LigB-specific regions, respectively. Values below each cartoon indicate the fold-regulation in the wild-type (WT) strain L495 parent compared to *perRA*, *perRB* and *perRA/B* mutant strains based on the corresponding RNA-Seq comparisons. **B.** Whole cell lysates of *L. interrogans* sv. Manilae strain L495 isogenic WT, *perRA*, *perRB*, *perRA/B* and *lvrAB* strains were generated from leptospires cultivated within DMCs, separated by SDS-PAGE, and probed with antiserum against repeats conserved in both LigA and LigB (gray region in panel A). After detection, membranes were stripped and re-probed using antiserum against recombinant FlaB1 as a loading control. Molecular weight markers (kDa) are shown on the left. Image is B is representative of results from three biological replicates, shown in S5 Fig.

The downregulated portion of the PerRA/B DMC regulon contains at least one gene potentially related to virulence. *LIMLP03665/colA*, encoding a collagenase precursor [137], was expressed at ~500-fold lower levels in the WT parent compared to the *perRA/B* mutant (Table 2). While collagenase-mediated degradation of host tissues likely enhances dissemination of leptospires during early infection [138], once in the kidneys, repression of *colA* could help reduce pathogen-mediated damage to renal epithelial cells. Further transcriptional analysis of this gene is needed to establish its expression profile in different tissues over the course of infection.

### Loss of LvrAB alone is not responsible for avirulence of the *perRA/B* double mutant in mice

As noted above, expression of *lvrAB* is disrupted only in the *perRA/B* double mutant (Table 1); similar results were obtained using leptospires grown *in vitro* [93]. Using LvrA- and LvrB-specific antisera, we confirmed our transcriptomic data at the protein level by immunoblot using whole cell lysates from WT, *perRA*, *perRB and perRA/B* mutant strains cultivated in DMCs (Figs 8A and S5A and S5C-D). Previously, Adhikarla *et al*. [19] reported that inactivation of *lvrAB* by transposon mutagenesis results in dysregulation of a large number of genes *in vitro,* including *ligB*. However, in our hands, we saw no decrease in LigA or LigB in the *lvrAB* mutant strain following cultivation in DMCs (Figs 7B and S5A-B). Adhikarla *et al*. [19] also reported that loss of either *lvrAB* or *lvrB* alone resulted in a significant loss of virulence in hamsters. To explore whether the avirulence of the *perRA/B* double mutant in mice (Fig 4C-E) is due solely to loss of LvrAB, we assessed the ability of *lvrAB* and *lvrB* transposon mutants to colonize the kidneys of C3H/HeJ mice (5 mice per strain, per experiment). While mice infected with either the *lvrAB* or *lvrB* mutant shed ~2-log_10_ less leptospires in their urine compared to the WT controls, all of the urine samples collected from mice infected with either mutant were darkfield positive by day 21 (Fig 8B). At day 28 p.i., kidneys harvested from all mice infected with either the *lvrAB* or *lvrB* mutant were positive for leptospires by both culturing in EMJH and qPCR (S6A Fig). Mice infected with the *lvrAB* mutant also seroconverted (S6B Fig). Thus, while LvrAB signal transduction contributes to virulence, loss of *lvrAB* expression alone is not responsible for the complete loss of virulence observed with the *perRA/B* double mutant in mice.

**Fig 8.**
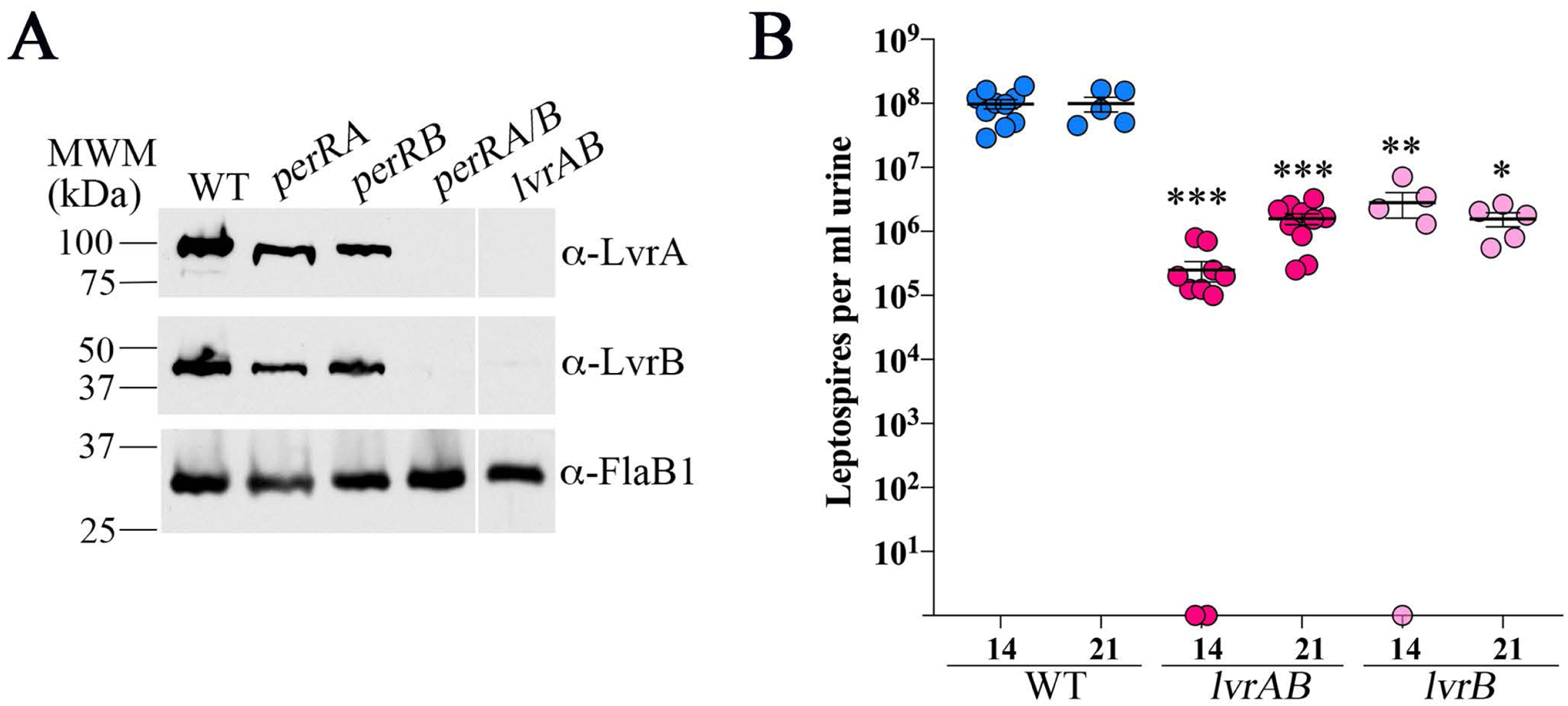
Expression of LvrAB requires at least one functional PerR homolog but the absence of LvrAB alone is not solely responsible for avirulence of the *perRA/B* double mutant. **A.** Whole cell lysates of *L. interrogans* sv. Manilae strain L495 wild-type (WT), *perRA*, *perRB*, *perRA/B* and *lvrAB* mutant strains were generated from leptospires cultivated within DMCs. Lysates were separated by SDS-PAGE, transferred to nitrocellulose, and probed with rabbit polyclonal LvrA- or LvrB-specific antiserum. Membranes were stripped and re-probed using rat polyclonal antiserum against recombinant FlaB1 as a loading control. Molecular weight markers (kDa) are shown on the left. **B.** Enumeration of leptospires in urine collected from C3H/HeJ mice 14- and 21-days post-infection following intraperitoneal inoculation with 10^5^ of wild-type (WT), *lvrAB* or *lvrB* mutant strains. Circles represent data for urine from individual mice (5 mice per group, per strain, per experiment). Bars represent the average and standard error of the mean. *p*-values were determined by comparing burdens in mice infected with wild-type (WT) and mutant strains at the same timepoint using a two-tailed *t*-test. ***, *p ≤* 0.0001; **, *p* = 0.0007; and *, *p* =0.0079.

## Discussion

*L. interrogans* must sense and respond to diverse signals and threats during the free-living and reservoir host phases of its zoonotic cycle. Not surprisingly, *L. interrogans* encodes substantially more sensory and regulatory proteins than *B. burgdorferi* and *T. pallidum* [139], two pathogenic spirochetes with far more restrictive growth niches. However, the regulatory networks and gene products that sustain *L. interrogans* in nature remain poorly understood. To gain insight into the transcriptomic changes that leptospires undergo within the host, we previously compared *L. interrogans* sv. Copenhageni strain Fiocruz L1-130 cultivated *in vitro* and in mammals using our DMC peritoneal implant model [41, 110]. From these studies emerged >100 genes that were differentially expressed in response to host-specific signals, including *LIC12034*, encoding the peroxide stress response regulator PerRA, which was upregulated 3.83-fold in DMCs. Herein, we confirmed these data using *L. interrogans* sv. Manilae strain L495 and also established that the three remaining FUR family regulators are transcribed at comparable (*perRB*) or higher (*ffr1* and *ffr2*) levels in DMCs compared to *in vitro*. The importance of FUR family regulators for host adaptation was confirmed recently by Zavala-Alvarado *et al*. [93], who demonstrated that leptospires lacking both PerRA and PerRB are unable to infect hamsters. In our current study, we establish that these regulators also are required for renal colonization of C3H/HeJ mice. In both animal models, loss of virulence was observed only when both PerRA and PerRB were inactivated, suggesting that these regulators may serve redundant or overlapping functions *in vivo*. Our finding that the *perRA*/*B* double mutant survives at wild-type levels in DMCs is particularly noteworthy as it demonstrates that the avirulent phenotype observed for this mutant is not due to a metabolic lesion (*i.e*., metal starvation) but instead reflects dysregulation of one or more virulence-related genes. Transcriptomic analyses of *perRA* and *perRB* single and double mutants cultivated in DMCs brought to light a number of novel aspects of FUR-mediated regulation in *L. interrogans*. Most notably, the majority of genes in the PerRA, PerRB and PerRA/B regulons were differentially expressed only in DMCs, highlighting the importance of mammalian host-specific signals for PerR-mediated regulation in *L. interrogans*. Remarkably, inactivation of both PerRA and PerRB resulted in a DMC regulon that differs substantially from those of either single mutant and includes a large cohort of genes involved in environmental sensing, signal transduction and transcriptional regulation.

Despite several attempts, Zavala-Alvarado *et al.* [93] was unable to restore virulence to the *perRA/B* <u>double</u> mutant by *trans*-complementation with *perRA* or *perRB* alone. Consequently. We cannot rule out the possibility that the loss of virulence observed with the *perRA/B* double mutant is due to a spontaneous genetic defect outside of *perRA* or *perRB*. However, by comparative genomic sequencing of WT, *perRA/B* and *perRB* (the parental background for the *perRA/B* double mutant) strains, Zavala-Alvarado *et al*. [93] identified only two differences in the *perRA/B* strain. The first was a single nucleotide insertion in LIMLP11570, encoding a putative 3-oxoacyl ACP synthase related fatty acid synthesis. It is important to note that this same insertion also is observed in several *L. interrogans* isolates from human and animals and, as noted earlier, we saw no difference in the growth of the single and double mutants either *in vitro* or in DMCs. The second nonsynonymous difference is in LIMLP01895, encoding a putative hybrid histidine kinase; the corresponding polymorphism results in an alanine to valine substitution at amino acid 146; the position of this mutation is within an inter-domain region and, therefore, not likely to affect the protein’s putative signal transduction function(s). Nonetheless, further investigation is necessary to establish the extent to which LIMLP01895 contributes to gene regulation and/or loss of virulence in the *perRA/B* double mutant.

The presence of multiple FUR family regulators in *Leptospira* spp. was noted previously by Louvel *et al*. [44], who identified five distinct orthologs between *L. interrogans* and *L. biflexa*. Phylogenetic analyses presented herein identified a sixth FUR family regulator and established that two (PerRA and Ffr1) are conserved within both pathogenic and saprophytic species and two each are unique to either pathogenic (PerRB and Ffr2) or saprophytic subclades (PerRC and Ffr3). As their designations suggest, based on sequence alignments, three are predicted to function as PerRs. Thus far, only *B. licheniformis* has been shown to encode multiple PerRs (PerR_BL_, PerR2 and PerR3), each of which displays a different level of sensitivity to H_2_O_2_ (PerR2 > PerR_BL_ > PerR3) [140]; the extent of regulatory overlap between these three PerRs has yet to be determined. The peroxide responsiveness and/or metal sensing properties of the remaining three leptospiral FUR family regulators cannot be predicted based on sequence alone. Our finding that almost all saprophytic and pathogenic *Leptospira* spp. encode closely-related PerRA and Ffr1 orthologs, however, implies that these two regulators could function outside of a host (*e.g*., within soil and/or water). The presence of a single PerR (PerRB) in all but one of the four P2 subclade species examined (*L. wolfii*) may contribute to the ‘intermediate’ virulence of these *Leptospira* spp. compared to highly virulent P1 subclade [6, 141].

*L. interrogans* cultivated in DMCs express increased levels of catalase, AhpC-type peroxiredoxin and cytochrome c peroxidase [41], three enzymes typically associated with detoxification of reactive oxygen species (ROS) in bacteria [85, 87, 142]. These data also provide strong evidence that *L. interrogans* is exposed to ROS *in vivo* [41]. Consistent with this notion, catalase-deficient leptospires are more susceptible to H_2_O_2_ *in vitro* and show reduced virulence in hamsters [143]. Host phagocytic cells, which generate oxygen radicals *via* a dedicated NADPH oxidase [144, 145], are one likely source of exogenously-derived ROS *in vivo*. Leptospires within renal tubules, a highly oxygenated niche, also would be exposed to elevated levels of oxygen. Incomplete reduction of oxygen by iron-containing cytochromes is another potential source of endogenous ROS [85, 142]. In bacteria, oxidative stress responses often are coordinated by two evolutionarily distinct master regulators -- OxyR and PerR. OxyR, the more common of the two, belongs to the LysR family and functions primarily as an activator [146]. In its oxidized state, OxyR activates transcription of genes involved in the detoxification of H_2_O_2_ (catalase and AhpC), the prevention or repair of DNA damage (Dps) and/or redox homeostasis (glutathione reductase, thioredoxin) [85]. PerR, first described in *B. subtilis* [147], typically represses rather than activates many of the same genes as OxyR and is released from DNA by peroxidation [49, 54, 90]. Although OxyR and PerR regulate transcription by different mechanisms, they react with H_2_O_2_ at essentially the same rate constant (10^5^ M^-1^ s^-1^) [89] and orchestrate highly similar responses. *L. interrogans* does not encode an OxyR homolog but, as noted above, encodes at least two PerR orthologs, PerRA and PerRB. Consistent with PerR functions in other bacteria, as shown here and elsewhere [91, 93, 94], *L. interrogans perRA* mutants show enhanced survival following exposure to lethal levels of H_2_O_2_ *in vitro* and increased expression levels of catalase, AhpC and cytochrome c peroxidase *in vitro*. While inactivation of *perRB* had no effect on the ability of leptospires to withstand killing by H_2_O_2_, the *perRB* mutant showed increased tolerance to the superoxide-generating compound paraquat [93]. Moreover, no genes associated with ROS defenses were dysregulated in the *perRB* mutant *in vitro* [93]. In DMCs, only cytochrome c peroxidase, AhpC and a glutaredoxin were dysregulated in the double mutant. Interestingly, all three genes were expressed at higher levels in the WT compared to the mutant, suggesting that they are activated rather than repressed by PerRB. Moreover, expression of catalase was not significantly different in the WT vs. *perRB* or *perRA/B* DMC comparison.

The above data argue that while PerRA and PerRB may be ‘activated’ by ROS, the adaptive responses they control likely extend beyond oxidative stress. The prototypical PerR in *B. subtilis* (PerR_Bs_) can coordinate either Mn^2+^ or Fe^2+^.When co-factored with Fe^2+^, DNA binding by PerR:Fe is highly sensitive to H_2_O_2_ due to irreversible iron-dependent oxidation of metal-coordinating histidine residues [54, 96]. When cofactored with Mn^2+^, however, PerR_Bs_ is able to bind DNA but is no longer peroxide sensitive [96]. In this way, PerR functions both as a peroxide responsive regulator and a ratiometric sensor for iron and manganese, altering its transcriptomic output based on intracellular metal availability and/or oxidative stress. Our finding that the PerRA/B DMC regulon contained only three genes related to oxidative stress and no genes related to iron homeostasis raises the possibility that PerRA and PerRB function *in vivo* may be regulated by metal availability rather than oxidative stress. Moreover, it is possible that both PerR:Fe and PerR:Mn regulate different cohorts of virulence genes, depending on the host milieu.

Consistent with differences in the putative DNA-binding helices, we saw very little overlap between the PerRA and PerRB DMC regulons. Seven of the eight genes common to both regulons are located in a single locus encoding a TonB-dependent transporter (TBDT) system. In Gram-negative bacteria, TBDT systems promote the uptake of substrates, such as iron siderophores, heme, vitamin B12, and carbohydrates, that are either poorly transported by non-specific outer membrane porins or are present in the extracellular milieu at low concentration [115, 148]. Substrate binding and uptake is mediated by high affinity, substrate-specific TonB-dependent receptor (TBDR) proteins, which form outer membrane-embedded 22-stranded *β*-barrels [115]. The energy required for substrate transport is provided in the form of proton motive force, which is transduced from the inner to outer membrane by the TonB-ExbB-ExbD complex [115]. *L. interrogans* encodes 11 putative TonB-dependent receptors and at least two complete TonB-ExbB-ExbD transporters. None of the leptospiral TBDRs possess a N-terminal extension capable of interacting with anti-sigma factors, similar to that of the iron and heme TBDRs FecA and HasR in *E. coli* and *Serratia marcescens*, respectively [149, 150]. Only one TBDT system, *LIMLP04240-04270*, is differentially regulated by PerRA and PerRB in DMCs. While the substrate(s) recognized by the TBDR (LIMLP04270) cannot be predicted based on sequence, mutagenesis studies on its ortholog in *L. biflexa* suggest that it is not essential for uptake of iron or heme *in vitro* [44]. Moreover, a *L. interrogans* transposon mutant containing an insertion in *LIMLP04270* is virulent in hamsters [93]. However, given the large number of TBDRs in *L interrogans*, one of these may compensate for loss of LIMLP04270 *in vitro* and/or *in vivo*. Interestingly, *in vitro*, expression of *LIMLP04240-04270* was dysregulated in the *perRA* and *perRA/B* mutants but not the *perRB* single mutant; only *LIMLP04255* was upregulated 1.93-fold in the WT compared to the *perRB* mutant [93]. A second TBDR, encoded by *LIMLP08410*, was repressed by PerRA/B only within DMCs. Together, these data suggest that mammalian host signals play a key role in modulating TonB-dependent nutrient uptake in *L. interrogans* and, moreover, that the activity of PerRB is enhanced *in vivo*.

By comparative RNA-Seq, we identified four distinct PerR regulatory categories in *L. interrogans* (Fig 9). The first two include genes whose expression is controlled exclusively by a single PerR (PerRA^only^ and PerRB^only^). The most straightforward explanation for this category is that PerRA and PerRB recognize different upstream boxes. Given that the *perRA* and *perRB* single mutants are fully virulent in hamsters [93] and mice, genes in these two categories either are not required in mammals or encode redundant functions. The third category, PerRA^and^B, includes the TBDT locus, described above, which requires PerRA and PerRB for expression. Presumably the upstream regions for PerRA^and^B loci contain separate PerRA- and PerRB-specific boxes, both of which must be engaged for transcription. The fourth category, PerRA^or^B, contains genes that are regulated by both PerRs but require only one for expression. The most likely explanation for this category is that the upstream regions for these genes contain separate PerRA and PerRB boxes, only one of which needs to be engaged for expression. Although less likely, PerRA and PerRB also could recognize a single ‘degenerate’ PerR box. None of these scenarios, however, explains why the majority of PerRA^only^ and PerRB^only^ genes continue to be expressed in the *perRA/B* mutant in DMCs. We hypothesize that this unexpected regulatory scheme reflects a natural requirement for these gene products at points during the zoonotic cycle when both PerRA and PerRB are inactive. We envision two non-mutually exclusive explanations for this intriguing finding: (i) The PerRA/B DMC regulon contains at least two putative DNA binding proteins (DBPs) that are downregulated by PerRA^or^B in wild-type leptospires. Continued expression of these DBPs in the *perRA/B* double mutant could help sustain expression of PerRA^only^ or PerRB^only^ genes. Alternatively, loss of PerRA *<u>and</u>* PerRB could lead to a physiological state that enables *L. interrogans*’ other FUR family regulators, Ffr1 and Ffr2, to “take over” expression of PerRA^only^ and PerRB^only^ genes. Examples of regulatory overlap between FURs in other bacteria are well documented [82, 151–155]. Further studies are needed to determine which, if any, of these scenarios are operative in *L. interrogans*.

**Fig 9.**
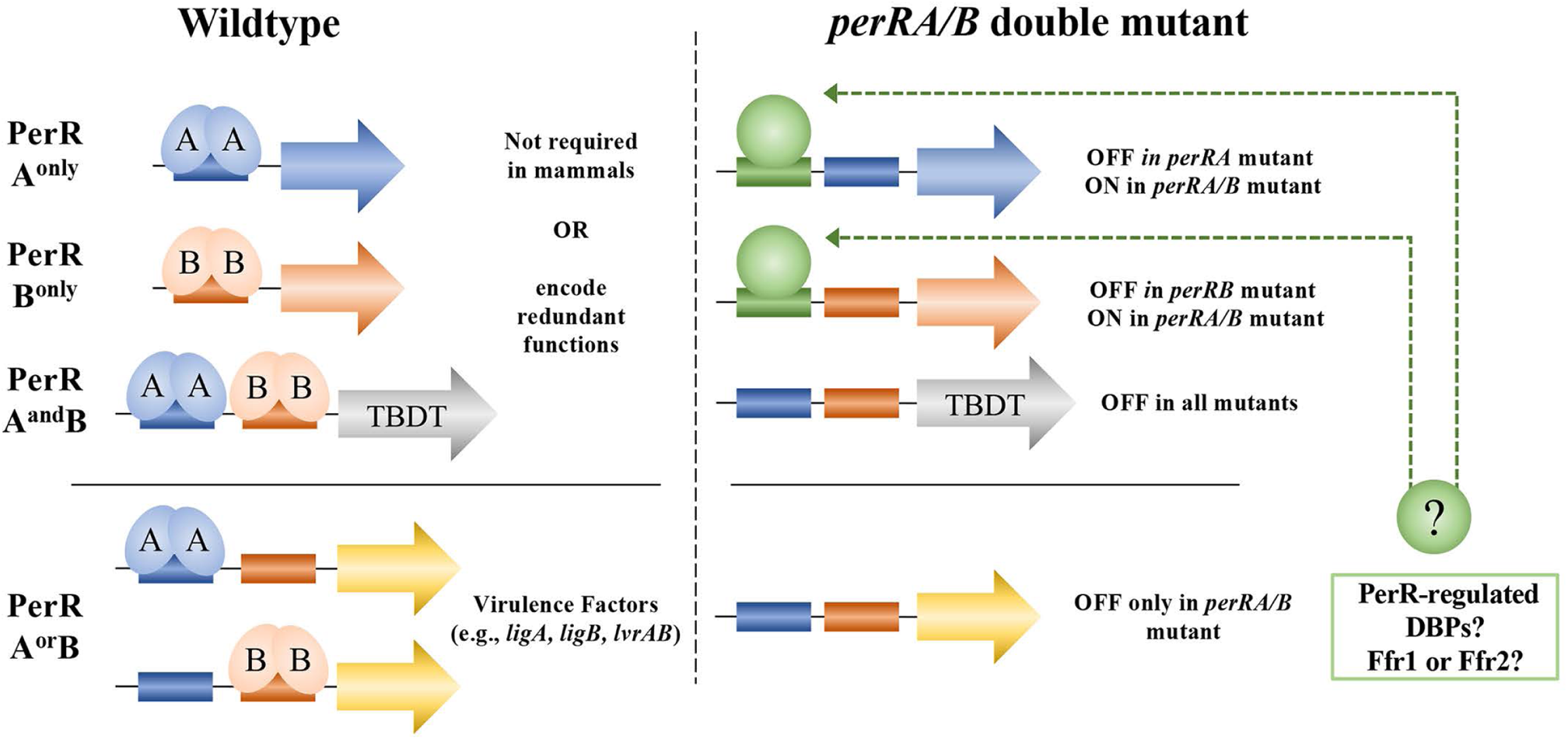
Working model to explain regulatory categories identified by RNA-Seq analyses of wild-type, *perRA*, *perRB* and *perRA/B* strains following cultivation in DMCs. Panels on the left and right indicate the expression profiles in wild-type and *perRA/B* strains. PerRA^only^, PerRB^only^, PerRA^and^B and PerRA^or^B categories are based on *wt* vs. *perRA wt* vs. *perRB,* and *wt* vs. *perRA/B* DMC regulons. DBPs, DNA binding proteins LIMLP19135 and LIMLP08975.

Surprisingly, the majority of genes controlled by PerRA and/or PerRB in DMCs were upregulated (*i.e*., expressed at lower levels in the single or double mutants compared to WT) rather than repressed *in vivo*. While only one prior study has demonstrated PerR-mediated activation [156], there are multiple examples of FUR family regulators acting as transcriptional activators [51, 52]. In *Vibrio vulnificus*, apo-Fur positively regulates its own expression by binding upstream of the *fur* promoter [157]. In *Helicobacter pylori* and *Salmonella enterica* sv. Typhimurium, Fur activates expression by binding upstream of target gene and helping to recruit RNA polymerase [158, 159]. In *α*-proteobacteria, Irrs (see below) act as positive and negative transcriptional regulators of genes related to heme homeostasis [59–62]. BosR, a FUR family regulator in the Lyme disease spirochete *B. burgdorferi*, activates transcription of the alternative sigma factor *rpoS* as part of a complex that includes the alternative sigma factor RpoN and the response regulator Rrp2 [160–165]. PerRA and/or PerRB also could activate transcription of target genes indirectly *via* repression of a regulatory small RNA (*e.g*., RyhB in *E. coli*) [166] or by preventing the binding of another repressor (*i.e*., anti-repression) [63, 64].

Designation of PerRA and PerRB as peroxide stress regulators in *L. interrogans* is based largely on *in vitro* studies showing increased survival of *perRA* and *perRB* mutants following exposure to H_2_O_2_ and paraquat, respectively [91, 93, 94]. Several lines of evidence, however, raise the possibility that these gene products function as iron response regulators (Irrs) rather than PerRs. Based on amino acid sequence alignments, PerRA and PerRB appear to be more closely related to Irrs than PerRs. In *α−*proteobacteria, Irrs and their regulatory partner, RirA, coordinate the expression of genes involved in heme biosynthesis with iron availability. Similar to FURs, RirA functions as metal-dependent transcriptional repressor but senses iron within Fe-S clusters rather than Fe^2+^. Interestingly, LIMLP06290 (LIC11283), annotated as a hypothetical protein, contains domains consistent with it being a RirA; the contribution of this putative RirA to iron homeostasis in *Leptospira* spp. has not been examined. At the sequence level, Irrs share a number of features with PerRs, including the presence of Asp and Arg residues in their regulatory metal sites and DNA binding helices, respectively. Irrs and PerRs also are responsive to similar levels of ROS, albeit by a different mechanism, and regulate many of the same effector genes (*i.e.,* catalases and peroxidases) [62, 167, 168]. In some, but not all cases, *irr* mutants also show increased survival *in vitro* under high H_2_O_2_ levels [62, 169]. As noted above, only *B. lichenformis* is known to encode multiple PerRs. Numerous bacteria, on the other hand, encode two or more Irrs [167]. Variable affinity of Irrs for their target promoters enables them to modulate gene expression over a wider range of conditions than PerRs [170]. The autoregulatory sequences identified upstream of *perRA* [92] diverge significantly from canonical PerR and Fur boxes but show strong similarity to “Irr-boxes” [171, 172]. Moreover, Irrs are known to act as activators as well as repressors [167, 171]. Although typically associated with peroxide-sensitive regulation of iron/heme acquisition and utilization, Irrs have been shown to control diverse cellular processes, including virulence. Moreover, the vast majority of histidine kinases upregulated by PerRA/B in DMCs contain one or more PAS-type sensor domains, which have been shown to function as heme sensors [173]. Given the established importance of heme for survival of *L. interrogans* in mammals [174–176], our findings raise the possibility that heme sensing by PerRA and/or PerRB in mammals could serve as an important initiating event for host adaptation.

Studies presented here and elsewhere [93] demonstrate for the first time that both PerRA and PerRB are required for full transcription of the virulence-related genes *ligA* and *ligB*. These pathogen-specific surface lipoproteins have been studied extensively for their contributions to host-pathogen interactions [177, 178], virulence [179] and potential use as vaccinogens [180–182]. Using a TALE-based transcriptional knockdown approach, Pappas and Picardeau [179] reported that both Ligs are required for virulence in hamsters. As noted earlier, *ligA* and *ligB* are not co-transcribed but instead share virtually identical upstream regions and, consequently, are co-regulated by the same environmental signals. Matsunaga, Haake and others previously reported that *ligA* and *ligB* are upregulated in response to physiological osmolarity (EMJH supplemented with 120 mM sodium chloride) [24] and increased temperature [183]. However, temperature-dependent regulation is mediated by a *cis*-acting RNA secondary structure that prevents translation at lower temperature and disruption of this *cis* element had minimal effect on osmoregulation [183]. Eshghi *et al*. [27] reported that inactivation of *lb139* (*LIMLP18410*), encoding a putative anti-ECF sigma factor, resulted in ~2.5-fold decreased expression of *ligB in vitro.* However, *LIMLP18410* is not in the PerRA/B regulon either *in vitro* [93] or in DMCs (this study) and, therefore, is not responsible for dysregulation of *ligA* and *ligB* in the *perRA/B* double mutant. Additional studies are needed to establish whether PerRA and/or PerRB regulate expression of *ligA* and *ligB* directly by binding to the *lig* promoter region or indirectly *via* another effector protein. The presence of multiple sensory and regulatory effector proteins in the PerRA and PerRB DMC regulons argues that activation of PerRA and PerRB, presumably by oxidative stress, initiates a complex regulatory network capable of sensing and responding to a wide range of mammalian host-specific signals. Our finding that LvrAB-deficient leptospires express normal levels of LigA and LigB argues that at least two PerRA/B-dependent regulatory pathways (LvrAB-dependent and -independent) are operative in *L. interrogans in vivo*.

## Material and Methods

### Ethics statement

All experiments involving animals conducted at UConn Health were performed in accordance with The Guide for the Care and Use of Laboratory Animals (8th Edition) (Guide for the Care and Use of Laboratory Animals, 1996) using protocols reviewed and approved by the UConn Health Institutional Animal Care and Use Committee [Animal Welfare Assurance (AWA) number A347-01].

### Bacterial cultivation *in vitro*

*L. interrogans* strains are described in S5 Table. Leptospires were cultivated routinely *in vitro* in Ellinghausen, McCullough, Johnson and Harris medium (EMJH) [184, 185] supplemented with 1% rabbit serum at 30°C under static conditions. Mutants were maintained in EMJH under appropriate antibiotic selection (spectinomycin, 40 μg/ml and/or kanamycin, 40 μg/ml). Cultures were harvested at late logarithmic phase (1-5 × 10^8^ per ml). Culture viability (*i.e.*, motility and cell morphology) was evaluated by darkfield microscopy. Leptospires were enumerated using a Petroff-Hausser counting chamber (Hausser Scientific Co., Horsham, PA). *Escherichia coli* strains were maintained in Lysogeny broth (LB) or LB agar supplemented with the appropriate antibiotics (ampicillin, 100 μg/ml; spectinomycin, 100 μg/ml; and/or kanamycin, 100 μg/ml). The genotypes of *L. interrogans* mutants used in these studies were confirmed by PCR and amplicon sequencing using primers listed in S6 Table.

### Routine DNA manipulation and cloning

Routine cloning was performed using In-Fusion HD Cloning Plus (Takara Bio USA Inc., Mountain View, CA) according to the manufacturer’s instructions. Plasmids were maintained in *E. coli* Top10 (Life Technologies, Grand Island, NY) or Stellar (TaKaRa, Mountain View, CA) cells and purified using QIAprep spin and midi kits (Qiagen, Valencia, CA). Bacterial genomic DNA was extracted from *L. interrogans* using the Gentra Puregene Yeast/Bacteria kit (Qiagen) according to the manufacturer’s recommendations. Routine and high-fidelity PCR amplifications were performed using RedTaq (Denville Scientific, Metuchen, NJ, United States) and CloneAmp HiFi (Takara Bio USA Inc., Mountain View, CA), respectively. DNA sequencing was performed by Genewiz, Inc. (Cambridge, MA). Routine sequence analyses were performed using MacVector (version 17.0.1, MacVector, Inc., Cary, NC, United States). Oligonucleotide primers used in these studies were purchased from Sigma-Aldrich (St. Louis, MO); primer sequences are provided in S6 Table.

### Generation of polyclonal antisera using recombinant protein

Recombinant His-tagged PerRA (LIMLP10155), PerRB (LIMLP05620), LvrA (LIMLP08490) and LvrB (LIMLP08485) cloned into pET28a vector (Novagen) and FlaB1 (LIMLP09410) cloned into pAE [186], were expressed in *E. coli* BL21 Star (DE3) or OverExpress C43 (DE3) (Lucigen/VWR, Radnor, PA). Following induction with IPTG, recombinant proteins were purified by nickel affinity chromatography using HisTrap Column (GE Healthcare Life Sciences Pittsburgh, PA). Rat polyclonal antisera against *L. interrogans* PerRA, PerRB, LvrA, LvrB and FlaB1 were generated by hyperimmunization of female Sprague-Dawley rats (Envigo, South Easton, MA) with 40-60 μg of recombinant His-tagged proteins co-administered with Freund’s Complete Adjuvant. After three weeks, two additional boosts of 40-60 μg of protein mixed 1:1 with Freund’s Incomplete Adjuvant were co-administered at two-week intervals. Two weeks after the second boost, animals were euthanized by anesthetic overdose and blood was collected by cardiac puncture. Sera was collected by centrifugation, aliquoted and frozen at −80°C.

### SDS–PAGE and immunoblot analyses

To analyze *L. interrogans* whole cell lysates, equivalent amounts of cells (~10^8^ leptospires per lane) were re-suspended and boiled in reducing Laemmli sample buffer (BioRad, Hercules, CA), separated through 10-12.5% separating polyacrylamide mini-gels and then visualized by GelCode Blue Stain Reagent (ThermoFisher Scientific, Grand Island, NY). Recombinant proteins expressed in *E. coli* were separated by SDS-PAGE and stained with GelCode Blue Stain Reagent (ThermoFisher). For immunoblotting, proteins were transferred to nitrocellulose membrane (GE Healthcare Life Sciences, Pittsburgh, PA) using Trans-Blot SD semi-dry transfer cell (BioRad, Hercules, CA). Membranes were blocked using milk block solution (MBS; 5% dry milk, 0.1% Tween 20, 5% fetal calf serum in PBS) for 1 h at room temperature. His-tagged recombinant proteins were detected using an HRP-conjugated anti-His monoclonal antibody (Sigma-Aldrich, St. Louis, MO) according to the manufacturer’s instructions. Antisera against recombinant His-tagged leptospiral proteins were diluted 1:500 (PerRA and PerRB), 1:1000 (LvrA, LvrB and FlaB1), 1:10,000 (LigA/B repeat region) in MBS and incubated overnight at 4°C. After washing with PBS containing 0.05% Tween 20 (PBST), bound antibody was detected with horseradish peroxidase-conjugated secondary antibody (Southern Biotechnology Associates, Birmingham, AL) diluted 1:30,000. After 1 hr at room temperature, membranes were washed at least five times with PBST and developed using the SuperSignal West Pico chemiluminescence substrate (Pierce, Rockford, IL).

### Generation of host-adapted leptospires

To obtain mammalian host-adapted organisms, *L. interrogans* sv. Manilae strain L495 wild-type and mutant strains were cultivated in DMCs as previously described [41, 42, 110]. Briefly, DMCs were prepared with 9-10 mls of EMJH medium (supplemented with an additional 10% bovine serum albumin to maintain osmotic pressure) at a starting inoculum of 10^4^ organisms per ml. Using strict aseptic technique, DMCs were implanted into the peritoneal cavity of an anesthetized female Sprague-Dawley rat. After nine days, animals were euthanized by CO2 narcosis and DMCs harvested. The viability and density of leptospires were evaluated by dark field microscopy using a Petroff-Hausser counting chamber (Hausser Scientific Co., Horsham, PA).

### Murine infection experiments

To determine the lethal dose to 50% of mice (LD_50_) for *L. interrogans* sv. Manilae strain L495, ten-week-old female C3H/HeJ mice (Jackson Laboratories, Bar Harbour, ME) were inoculated intraperitoneally (IP) with 200 μl of EMJH containing 5 × 10^6^, 10^6^, 10^5^ or 10^4^ leptospires (5 mice per group). Animals were monitored twice a day for signs of leptospirosis and, when moribund, were euthanized by anesthetic overdose. LD_50_ was calculated using the Reed-Muench method [187]. For virulence studies, 10^5^ of wild-type parent, mutant or complemented strains were used to infect C3H/HeJ mice (5 animals per group, per experiment). Beginning 14 days post-infection (p.i.), animals were monitored for the presence of leptospires in urine, collected in a metabolic chamber for ~45 min following subcutaneous administration of furosemide (2-10 mg/kg, IP). Burdens in urine were assessed by darkfield microcopy using a Petroff-Hausser counting chamber. Twenty-eight or 42 days p.i. (virulence and LD_50_ experiments, respectively), animals were euthanized by CO_2_ narcosis and blood and kidneys were collected for serology, culturing in EMJH, and qPCR. Sera from individual mice were used to probe whole cell lysates (~10^8^ leptospires per lane) prepared from the wild-type parent grown *in vitro* in EMJH at 30°C.

### qRT-PCR

Total RNA was isolated from leptospires (four biological replicates per condition) cultivated *in vitro* at 30°C or following cultivation in DMCs as previously described [41]. cDNAs (+ and – RT) were assayed in quadruplicate in 25 μl reactions performed with SsoAdvanced Universal SYBR or Probe (*lipL32*) Super Mixes (Bio-Rad). Oligonucleotide primers used for qRT-PCR are provided in S6 Table. Copy numbers were calculated using internal standard curves (10^7^ – 10^1^ copies) generated using purified amplicons for *perRA*, *perRB*, *LIMLP18590* and *LIMLP04825* and then normalized against *lipL32* [180]. The standard curve for *lipL32* was generated using a copy of the *lipL32* amplicon cloned into pCR2.1-TOPO plasmid (Invitrogen). Normalized copy numbers were compared using an unpaired *t* test with two-tailed *p* values and 95% confidence interval (Prism v. 6, GraphPad Software).

### Quantitation of burdens by qPCR

DNA was extracted from infected kidneys using the Qiagen DNeasy Blood & Tissue kit according to the manufacturer’s recommendations. DNAs were analyzed by quantitative PCR (qPCR) using a TaqMan-based assay for *lipL32* [180] in 25 μl reactions performed with SsoAdvanced Universal Probes Super Mix (Bio-Rad). Copy numbers for *lipL32* were determined using an internal standard curve for the *lipL32* amplicon cloned into pCR2.1 TOPO (Invitrogen). Average values for each strain were compared using an unpaired *t* test with two-tailed *p* values and 95% confidence interval (Prism v. 6, GraphPad Software).

### RNA sequencing and comparative transcriptomics

Total RNA was prepared from leptospires cultivated in DMCs using TRIzol Reagent (ThermoFisher) (3 biological replicates per strain) and then treated twice with TURBO DNase (ThermoFisher) followed by purification using RNeasy columns (Qiagen) as previously described [41]. Samples were eluted in RNAse-free water and purified RNA was analyzed using Qubit RNA HS Assay Kit (Thermo) and Agilent TapeStation 4200 (Agilent Technologies, Santa Clara, CA, USA) using the RNA High Sensitivity assay. Only samples with Ribosomal Integrity number (RIN*e*) values >7.5 were used for library preparation. Stranded libraries were prepared from ribo-depleted RNA using Zymo-Seq RiboFree Total RNA Library Kit according to manufacturer’s instructions. Libraries were validated for length and adapter dimer removal using the Agilent TapeStation 4200 D1000 high-sensitivity assay and then quantified and normalized using the double-stranded DNA (dsDNA) high-sensitivity assay for Qubit 3.0 (Life Technologies, Carlsbad, CA). Libraries were run on an Illumina High Output 75-cycle v2.5 NextSeq 500 flow cell. Raw reads for each sample were trimmed using Sickle (v. 1.3.3; available from https://github.com/najoshi/sickle) and then mapped using EDGE-pro version 1.1.3 [113] using fasta, protein translation table (ptt) and ribosomal/transfer RNA table (rnt) files based on the *L. interrogans* sv. Manilae strain UP-MMC-NIID LP genome (NZ_CP011931.1, NZ_CP011932.1 and NZ_CP011933.1). Differential expression was determined using DESeq2 [114]. Genes expressed at ≥3-fold higher/lower levels in the mutant compared to the wild-type parent with a False Discovery Rate (FDR)-adjusted *p*-value (*q*-value) ≤ 0.05 were considered differentially expressed. Raw read data have been deposited in the NCBI Sequence Read Archive (SRA) database (BioProject accession PRJNA659512, samples SRR12604412, SRR12604413, SRR12604414, SRR12604415, SRR12604416, SRR12604417, SRR12604418, SRR12604419, SRR12604420, SRR12604421, SRR12604422 and SRR12604423).

### Bioinformatics

Routine and comparative sequence analyses were performed using MacVector (version 17.5.4; MacVector, Inc., Apex, NC). Clusters of Orthologous Group (COG) classifications are based on MicroScope, an integrated platform for the annotation of bacterial gene function through genomic, pangenomic and metabolic comparative analysis [188]. Conserved domain searches were performed using Conserved Domain Database (CDD) Search [189], UniProt [190] and InterPro [191]. Candidate lipoproteins were identified based on Setubal *et al*. [192] and LipoP server [193]. Subcellular localization predictions were performed by BUSCA (Bologna Unified Subcellular Component Annotator) [194]. Multiple sequence alignments were generated by Clustal Omega [195] and MAFFT 7 [196]. Phylogenetic trees were generated using PhyML 3.0 [197] with LG substitution model chosen after an Akaike Information Criterion (AIC) model selection [198]. Tree improvement was done by subtree pruning and regrafting (SPR) method [199] with ten random starting trees. Robustness of branches was assessed by Approximate Likelihood-Ratio Test (aLRT-SH) [200]. The resulting trees were visualized and annotated using Interactive Tree of Life (iTOL, v 4.3) [201]. FUR domain-containing proteins in 26 *Leptospira* spp. genomes (10, 5, 6 and 5 species from subclade P1, P2, S1 and S2, respectively [98]) were identified using the *Leptospira* species name as a query in the Ferric-uptake regulator domain entry (IPR002481) in EMBL-EBI InterProScan [202]. Orthologs shared between *L. interrogans* sv. Manilae strain L495 and sv. Copenhageni strain Fiocruz L1-130 strains were identified using OrthoVenn 2.0 [203].

## Acknowledgements

The authors wish to thank James Matsunaga and David Haake (UCLA) for generously providing LigA/B antisera. We also would like to thank Elsio Wunder (Yale) providing *lvrAB* and *lvrB* mutants. We would like to acknowledge Melissa McLain for her technical assistance on these studies and Jessica Grassmann for her assistance generating figures. We are indebted to Bo Reese, Center for Genome Innovation (CGI; UConn Storrs), for her superb technical assistance with our RNA-Seq studies. Lastly, we extend our sincere thanks to Justin Radolf for helpful suggestions throughout all stages of this work and for his careful reading of the manuscript.

## Supporting Information

**S1 Fig. Multiple sequence alignment of FUR domain-containing proteins in representative *Leptospira* spp. from Fig 2**. Species from pathogenic subclades P1 and P2 are colored in black and blue, respectively, while saprophytic species from subclades S1 and S2 are in red and green, respectively. Genomic locus tags for each FUR family proteins in *Leptospira* spp. are indicated. Highly conserved residues predicted as regulatory metal binding sites are highlighted in yellow, green and magenta. Putative structural metal binding sites are highlighted in cyan. *Leptospira* species are abbreviated as follow: L. int, *L. interrogans*; L. kir, *L. kirschneri*; L. adl, *L. adleri*; L. als, *L. alstonii*; L. san, *L. santarosai*; L. bor, *L. borgpetersoni*; L. ale, *L. alexanderi*; L. wol, *L. wolfii*; L. lis, *L*. *liscerasiae*; L. ina, *L. inadai*; L. fai, *L. fainei*; L. bif, *L. biflexa*; L. mey, *L. meyeri*; L. ter, *L. terpstrae*; L. van, *L. vanthielii*; L. ryu, *L. ryugenii*; L. ily, *L. ilyithenensis*; L. ido, *L. idonii*.

**S2 Fig. Expression of PerRA and PerRB in *L. interrogans* wild-type and mutant strains**. Whole cell lysates of *L. interrogans* sv. Manilae strain L495 wild-type (WT), *perRA*, *perRB* and *perRA/B* strains cultivated *in vitro*. Lysates were separated by SDS-PAGE, transferred to nitrocellulose, and probed with rat polyclonal PerRA- or PerRB-specific antiserum. Membranes were stripped and re-probed using rat polyclonal antiserum against recombinant FlaB1 as a loading control.

**S3 Fig. PerRA and PerRB regulate the expression of a locus (*LIMLP04285-04240*) that includes a TonB-dependent transport system.** Data from comparative RNA-Seq analysis of wild-type (WT) vs. *perRA*, *perRB* and *perRA/B* strains identified a nine gene chromosomal locus that includes *lipL48* and genes encoding a TonB-dependent receptor and ExbB/ExbD/TonB transporter. Fold-of-regulation for each gene are based on RNA-Seq data from wild-type and mutant leptospires grown in DMCs, presented in S2-S4 Tables, and *in vitro* in EMJH at 30°C (IV), presented in Zavala-Alvarado *et al.* [93, 94]

**S4 Fig. Overview of genes differentially expressed by *L. interrogans perRA* and *perRB* single mutants.** Cluster of Orthologous Genes (COG) categorization of differentially expressed genes (DEGs) in the wild-type (WT) vs. *perRA* (**A**) and *perRB* (**B**) RNA-Seq comparisons. COG predictions for individual genes are presented in S2-S3 Tables. Number of DEGs in each COG are indicated on the *x*-axis.

**S5 Fig. Expression of LigA, LigB, LvrA and LvrB are reduced in *perRA* and *perRB* single mutants and undetectable in the *perRA/B* double mutant. A.** Whole cell lysates of *L. interrogans* sv. Manilae strain L495 isogenic wildtype (WT), *perRA*, *perRB*, *perRA/B* and *lvrAB* strains were generated from leptospires cultivated within DMCs, separated by SDS-PAGE, probed with polyclonal antiserum against LvrA, LvrB, or N-terminal conserved repeat region for LigA/LigB. Panels represent independent biological replicates and detected by chemiluminescence imaging as described in Methods. After detection, membranes were stripped and re-probed using polyclonal antiserum against recombinant FlaB1 as a loading control. Intensity values for LigA/B (combined), LvrA, and LvrB in each replicate were quantified using ImageJ and the normalized based on values for FlaB1 in the same lysate. Normalized values for mutant strains were compared to those from the WT, which was set to 100. Bars represent the standard error of the mean from three biological replicates. Significant was determined in Prism (GraphPad) using a two-tailed *t*-test. Different letters indicate a significant difference (*p ≤* 0.05) in pairwise comparisons.

**S6 Fig. Expression of LvrAB requires at least one functional PerR homolog but the absence of LvrAB alone is not solely responsible for avirulence of the *perRA/B* double mutant. A.** Burdens of leptospires in kidneys harvested from mice in Fig 8B. DNA samples from kidneys harvested 28 days post-inoculation were assessed (in quadruplicate) by qPCR using a Taqman-based assay for *lipL32*. Bars represent the average and standard error of the mean. *p*-values were determined by comparing burdens in mice infected with wild-type (WT) and mutant strains at the same timepoint using a two-tailed *t*-test; we saw no significant difference (*p*>0.05) between burdens between the WT, *lvrAB* and *lvrB* strains. **B**. Immunoblot analysis of sera collected from C3H/HeJ mice 28-days following intraperitoneal inoculation with 10^5^ wild-type or *lvrAB* mutant strains and then used to probe whole cell lysates of *L. interrogans* sv. Manilae strain L495 grown in EMJH at 30°C.

**S1 Table. Summary of RNA-Seq raw read data.**

**S2 Table. Comparative RNA-Seq data for *L. interrogans* sv. Manilae L495 wild-type and *perRA* strains cultivated in dialysis membrane chambers (DMCs)**. The genome sequence of *L. interrogans* sv. Manilae strain UP-MMC-NIID LP (accession numbers NZ_CP011931.1, NZ_CP011932.1 and NZ_CP011933.1) was used for mapping and differential gene expression analysis. All non-coding RNAs and pseudogenes were removed before DESeq2 analysis. **Column A**: RefSeq locus tag. **Column B**: Locus tag. **Column C**: *L. interrogans* serovar Copenhageni strain Fiocruz L1-130 (accession numbers NC_005823.1 and NC_005824.1) orthologs identified using OrthoVenn 2.0 [203]. Dashes (-) indicate genes for which no clear ortholog was identified. **Column D**: Description of gene product, following genome annotation. **Columns E, F**: Clusters of Orthologous Group (COG) classifications based on MicroScope [188]. **Column G**: Fold-regulation of the corresponding gene based on RNA-Seq analysis by Zavala-Alvarado *et al.* [93, 94] performed using the same strains cultivated *in vitro*. Positive and negative numbers indicate upregulation and repression, respectively, in the WT compared to the mutant strain. Dashes (-) indicate genes that are not differentially regulated at least 3-fold (*p*≤0.05) by PerRA *in vitro*. **Column H**: Fold-regulation determined using DESeq2 based on WT vs. *perRA* mutant RNA-Seq analysis using leptospires cultivated within DMCs. **Column I**: Type of regulation by PerRA in DMCs. Genes expressed at *≥*3-fold higher/lower levels in the WT vs. mutant with a False-discovery rate-adjusted-*p* value (*q*) *<u>≥</u>*0.05 were considered differentially expressed. “NO” indicates genes that are not regulated by PerRA in DMCs; “Up” indicates genes upregulated by PerRA in DMCs (expressed at lower levels in the mutant vs. WT); “Down” indicates genes downregulated by PerRA in DMCs (expressed at higher levels in the mutant vs WT). **Columns J-O**: Number of mapped reads per gene for each one of the three biological replicates per strain. **Columns P-AC**: Output from DESeq2 for WT vs. *perRA* mutant strains cultivated in DMCs (3 biological replicates per strain). **Column P**: Mean DESeq2 values for each gene. **Column Q**: Log_2_-fold change in gene expression. **Column R**: Power function transformation of log_2_-fold change. **Column S**: Fold regulation. **Column T-W**. Statistical analysis of differential gene expression including standard error estimate for the log_2_-fold change estimate (lfcSE, column T) and adjusted *p*-value (W). **Columns X-AC**: Normalized copy numbers per gene (3 biological replicates per strain).

**S3 Table. Comparative RNA-Seq data for *L. interrogans* sv. Manilae L495 wild-type and *perRB* strains cultivated in dialysis membrane chambers (DMCs)**. The genome sequence of *L. interrogans* sv. Manilae strain UP-MMC-NIID LP (accession numbers NZ_CP011931.1, NZ_CP011932.1 and NZ_CP011933.1) was used for mapping and differential gene expression analysis. All non-coding RNAs and pseudogenes were removed before DESeq2 analysis. **Column A**: RefSeq locus tag. **Column B**: Locus tag. **Column C**: *L. interrogans* serovar Copenhageni strain Fiocruz L1-130 (accession numbers NC_005823.1 and NC_005824.1) orthologs identified using OrthoVenn 2.0 [203]. Dashes (-) indicate genes for which no clear ortholog was identified. **Column D**: Description of gene product, following genome annotation. **Columns E, F**: Clusters of Orthologous Group (COG) classifications based on MicroScope [188]. **Column G**: Regulation of the corresponding gene based on RNA-Seq analysis by Zavala-Alvarado *et al.* [93, 94] performed using the same strains cultivated *in vitro*. Positive and negative numbers indicate upregulation and repression, respectively, in the WT compared to the mutant strain. Dashes (-) indicate genes that are not differentially regulated at least 3-fold (*p*≤0.05) by PerRB *in vitro*. **Column H**: Fold-regulation determined using DESeq2 based on WT vs. *perRB* mutant RNA-Seq analysis using leptospires cultivated within DMCs. **Column I**: Type of regulation by PerRB in DMCs. Genes expressed at *≥*3-fold higher/lower levels in the WT vs. mutant with a False-discovery rate-adjusted-*p* value (*q*) *<u>≤</u>* 0.05 were considered differentially expressed. “NO” indicates genes that are not regulated by PerRB in DMCs; “Up” indicates genes upregulated by PerRB in DMCs (expressed at lower levels in the mutant vs. WT); “Down” indicates genes downregulated by PerRB in DMCs (expressed at higher levels in the mutant vs. WT). **Columns J-O**: Number of mapped reads per gene for each one of the three biological replicates per strain. **Columns P-AC**: Output from DESeq2 for WT vs. *perRB* mutant strains cultivated in DMCs (3 biological replicates per strain). **Column P**: Mean DESeq2 values for each gene. **Column Q**: Log_2_ fold change in gene expression. **Column R**: Power function transformation of log_2_-fold change. **Column S**: Fold regulation. **Column T-W**. Statistical analysis of differential gene expression including standard error estimate for the log_2_-fold change estimate (lfcSE, column T) and adjusted *p*-value (W). **Columns X-AC**: Normalized copy numbers per gene (3 biological replicates per strain).

**S4 Table. Comparative RNA-Seq data for *L. interrogans* sv. Manilae L495 wild-type and *perRA/B* strains cultivated in dialysis membrane chambers (DMCs)**. The genome sequence of *L. interrogans* sv. Manilae strain UP-MMC-NIID LP (accession numbers NZ_CP011931.1, NZ_CP011932.1 and NZ_CP011933.1) was used for mapping and differential gene expression analysis. All non-coding RNAs and pseudogenes were removed before DESeq2 analysis. **Column A**: RefSeq locus tag. **Column B**: Locus tag. **Column C**: *L. interrogans* sv. Copenhageni strain Fiocruz L1-130 (accession numbers NC_005823.1 and NC_005824.1) orthologs identified using OrthoVenn 2.0. [203]. Dashes (-) indicate genes for which no clear ortholog was identified. **Column D**: Description of gene product, following genome annotation. **Column E**. Identification of conserved domain(s) within the corresponding gene product based on search of the Interpro database [191, 202]. The domain identification for each gene is followed by description; [D] indicates a domain; [F] indicates a protein family; [H] indicates a homologous superfamily. **Column F**. Uniprot entry for the orthologous gene in *L. interrogans* sv. Copenhageni strain Fiocruz L1-130 genome. **Columns G, H**: Clusters of Orthologous Group (COG) classifications based on MicroScope [188]. **Column I**: Regulation of the corresponding gene based on RNA-Seq analysis by Zavala-Alvarado *et al.* [93, 94] performed using the same strains cultivated *in vitro*. Positive and negative numbers indicate upregulation and repression, respectively, in the WT compared to mutant strain. Dashes (-) indicate genes that are not differentially regulated at least 3-fold (*p≤*0.05) by PerRA/B *in vitro*. **Column J**: Fold-regulation determined using DESeq2 based on WT vs. the *perRA/B* double mutant RNA-Seq analysis using leptospires cultivated within DMCs. **Column K**: Type of regulation by PerRA/B in DMCs. Genes expressed at *≥*3-fold higher/lower levels in the WT versus mutant with a False-discovery rate-adjusted-*p* value (*q*) ≤0.05 were considered differentially expressed. “NO” indicates genes that are not regulated by PerRA/B in DMCs; “Up” indicates genes upregulated by PerRA/B in DMCs (expressed at lower levels in the mutant vs. WT); “Down” indicates genes downregulated by PerRA/B in DMCs (expressed at higher levels in the mutant vs. WT). **Column L**: Behavior of the corresponding gene in the WT vs. *perRA* (A) and *perRB* (B) mutants by RNA-Seq using *L. interrogans* cultivated in DMCs. “Up” and “Down”, respectively, are used to significant (*≥*3-fold, *q≤*0.05) upregulation or repression of the gene in the WT compared to the mutant. **Columns M-R**: Number of mapped reads per gene for each one of the three biological replicates per strain. **Columns S-AF**: Output from DESeq2 for WT vs. *perRA/B* mutant strains cultivated in DMCs (3 biological replicates per strain). **Column S**: Mean DESeq2 values for each gene. **Column T**: Log_2_ fold change in gene expression. **Column U**: Power function transformation of Log_2_ fold change. **Column V**: Fold regulation. **Column W-Z**. Statistical analysis of differential gene expression including standard error estimate for the log2 fold change estimate (lfcSE, column T) and adjusted *p*-value (W). **Columns AA-AF**: Normalized copy numbers per gene (3 biological replicates per strain).

**Table S5. Bacterial strains used in these studies.**

**Table S6. Oligonucleotide primers used in these studies.**

## Notes

### Competing Interest Statement

The authors have declared no competing interest.

