## Supplementary Information for "The FUR-like regulators PerRA and PerRB integrate a complex regulatory network that promotes mammalian host-adaptation and virulence of *Leptospira interrogans*"

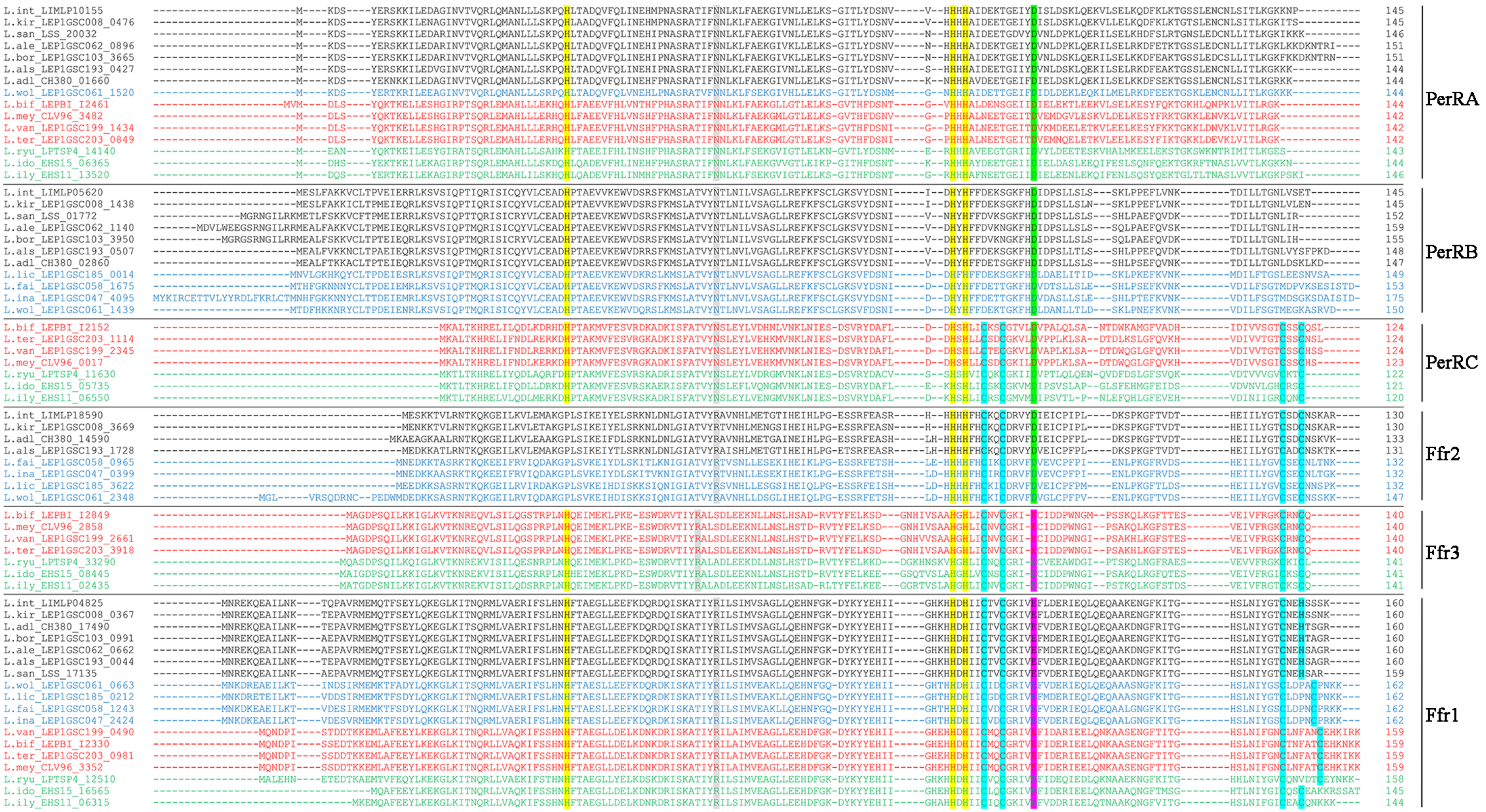

**S1 Figure. Multiple sequence alignment of FUR domain-containing proteins in representative *Leptospira* spp. from Figure 2.** Species from pathogenic subclades P1 and P2 are colored in black and blue, respectively, while saprophytic species from subclades S1 and S2 are in red and green, respectively. Genomic locus tags for each FUR family proteins in *Leptospira* spp. are indicated. Highly conserved residues predicted as regulatory metal binding sites are highlighted in yellow, green and magenta. Putative structural metal binding sites are highlighted in cyan. *Leptospira* species are abbreviated as follow: L. int, *L. interrogans*; L. kir, *L. kirschneri*; L. adl, *L. adleri*; L. als, *L. alstonii*; L. san, *L. santarosai*; L. bor, *L. borgpetersoni*; L. ale, *L. alexanderi*; L. wol, *L. wolfii*; L. lis, *L. liscerasiae*; L. ina, *L. inadai*; L. fai, *L. fainei*; L. bif, *L. biflexa*; L. mey, *L. meyeri*; L. ter,

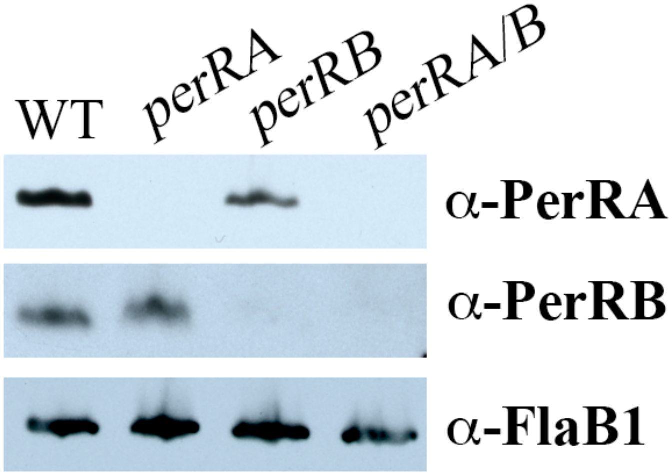

**S2 Figure. Expression of PerRA and PerRB in *L. interrogans* wild-type and mutant strains**. Whole cell lysates of *L. interrogans* sv. Manilae strain L495 wild-type (WT), *perRA*, *perRB* and *perRA/B* strains cultivated *in vitro*. Lysates were separated by SDS-PAGE, transferred to nitrocellulose, and probed with rat polyclonal PerRA- or PerRB-specific antiserum. Membranes were stripped and reprobed using rat polyclonal antiserum against recombinant FlaB1 as a loading control.

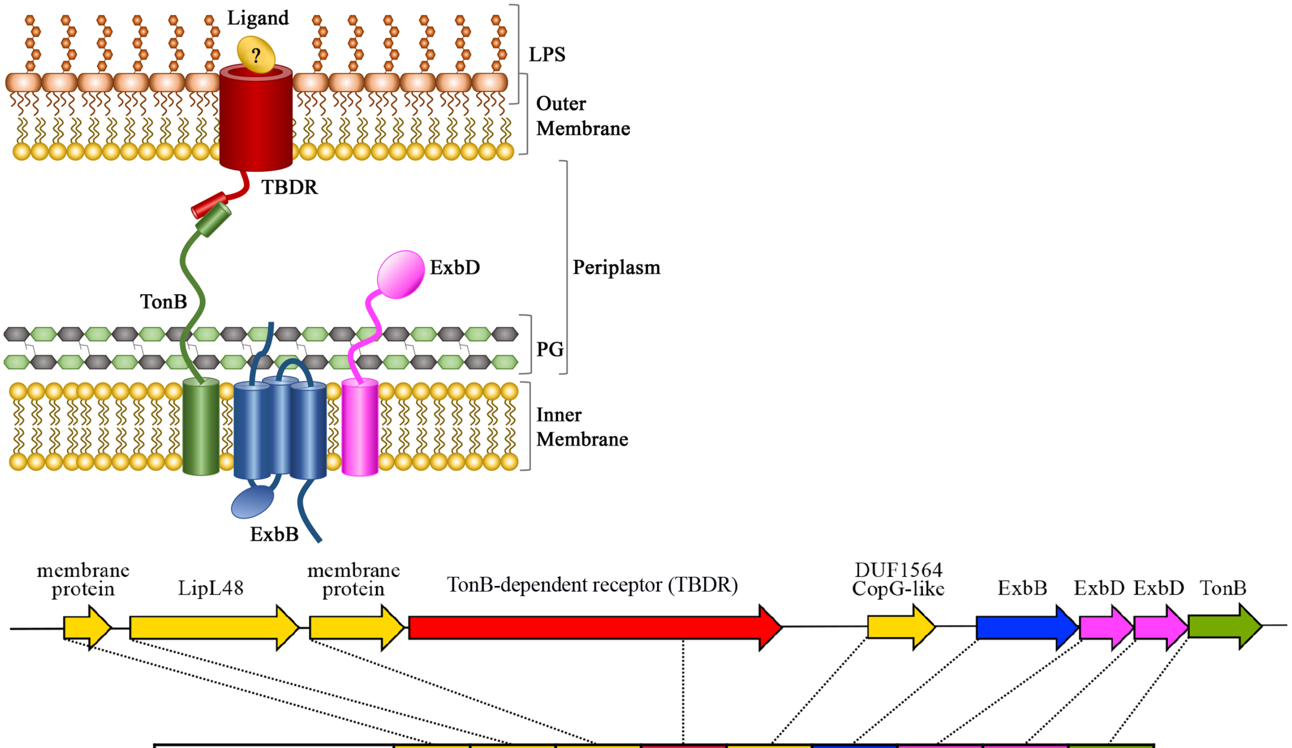

| WT vs. Mut (RNA-Seq ) | | 04285 | 04280 | 04275 | 04270 | 04260 | 04255 | 04250 | 04245 | 04240 |
|---|---|---|---|---|---|---|---|---|---|---|
| *perRA* | DMC | 5.4 | 170.3 | 23.0 | 35.4 | - | 28.1 | 22.9 | 11.3 | 8.6 |
| | IV | - | 45.4 | 14.8 | 9.59 | - | 44.6 | 40.9 | 24.3 | 24.3 |
| *perRB* | DMC | 3.0 | 8.0 | 13.8 | 5.6 | 6.1 | - | 5.3 | 10.3 | 4.1 |
| | IV | - | - | - | - | - | - | - | - | - |
| *perRA/B* | DMC | - | 4.9 | 6.7 | 3.7 | - | - | 3.3 | 3.8 | 3.2 |
| | IV | 6.4 | 5.9 | 5.7 | - | 3.2 | 5.9 | 5.0 | 4.9 | 4.2 |

**S3 Figure. PerRA and PerRB regulate the expression of a locus (*LIMLP04285-04240*) that includes a TonB-dependent transporter system (TBDT).** Data from comparative RNA-Seq analysis of wild-type (WT) vs. *perRA*, *perRB* and *perRA/B* strains identified a nine gene chromosomal locus that includes *lipL48* and genes encoding a TonB-dependent receptor and ExbB/ExbD/TonB transporter. Fold-of-regulation for each gene are based on RNA-Seq data from WT and mutant leptospires grown in DMCs, presented in Tables S2-S4, and *in vitro* in EMJH at 30°C (IV), presented in Zavala-Alvarado *et al.* [1, 2]

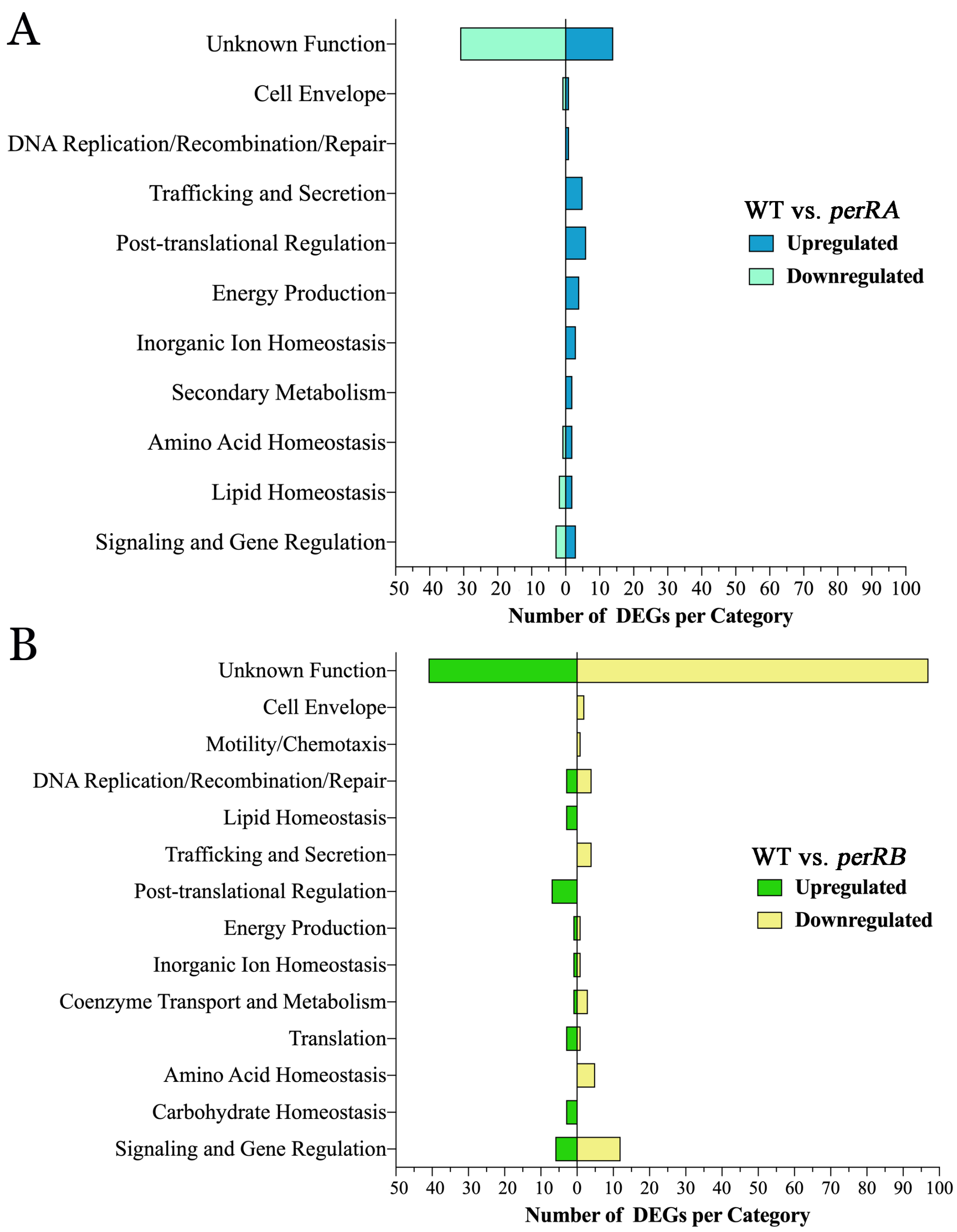

**S4 Figure. Overview of genes differentially expressed by *L. interrogans perRA* and *perRB* single mutants.** Cluster of Orthologous Genes (COG) categorization of differentially expressed genes (DEGs) in the wild-type vs. *perRA* (**A**) and *perRB* (**B**) RNA-Seq comparisons. COG predictions for individual genes are presented in Tables S2-S3. Number of DEGs in each COG are indicated on the *x*-axis.

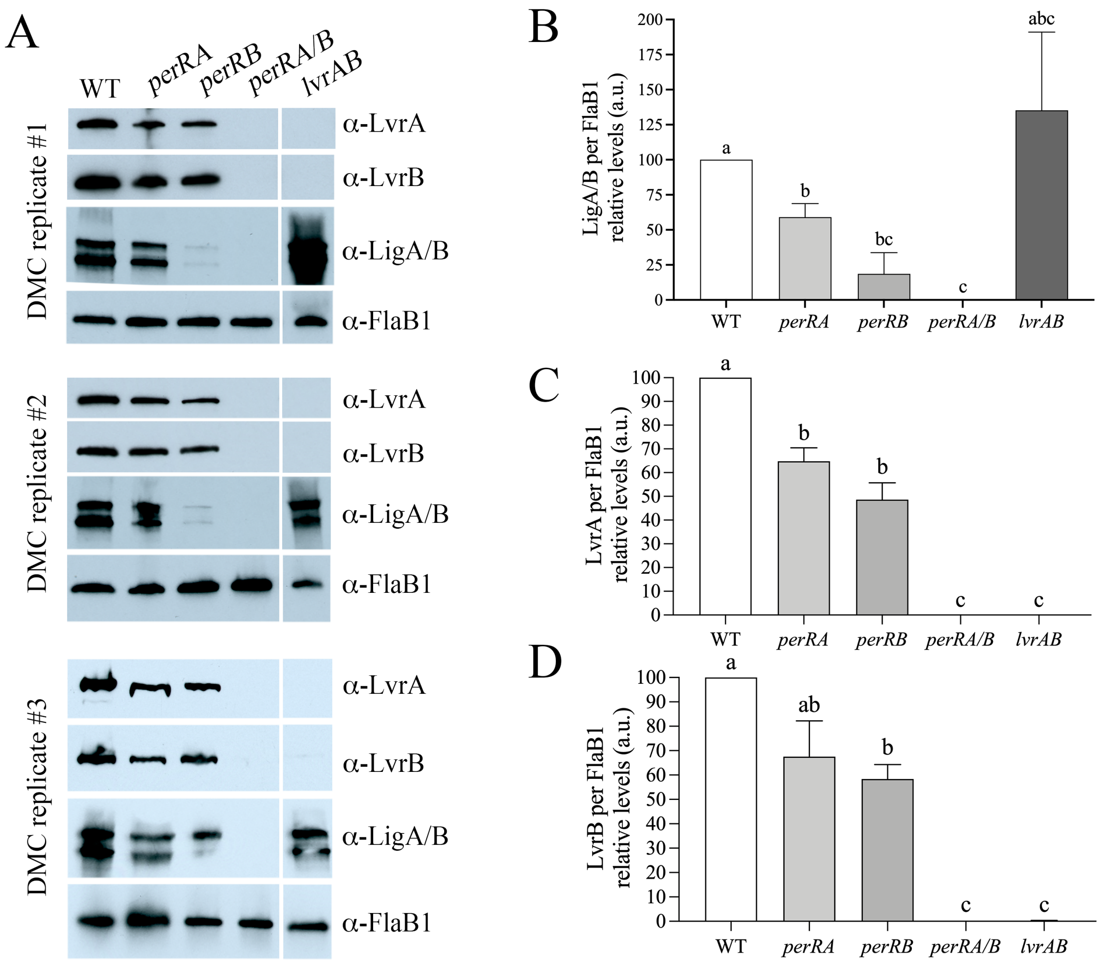

**S5 Figure. Expression of LigA, LigB, LvrA and LvrB is reduced in *perRA* and *perRB* single mutants and undetectable in the *perRA/B* double mutant. A.** Whole cell lysates of *L. interrogans* sv. Manilae strain L495 isogenic wild-type (WT), *perRA*, *perRB*, *perRA/B* and *lvrAB* strains were generated from leptospires cultivated within DMCs, separated by SDS-PAGE, probed with antiserum against LvrA, LvrB, or N-terminal conserved repeat region for LigA/LigB. Panels in **A** represent independent biological DMCs. After detection, membranes were stripped and re-probed using antiserum against FlaB1 as a loading control. **B-D.** Intensity values for LigA/B (combined), LvrA, and LvrB in each replicate were quantified using ImageJ and the normalized based on values for FlaB1 in the same lysate. Normalized values for mutant strains were compared to those from the WT, which was set to 100. Bars represent the standard error of the mean from three biological replicates. Significance was determined using a two-tailed *t*-test in Prism (GraphPad). Different letters indicate a significant difference ($p \leq 0.05$) in pairwise comparisons.

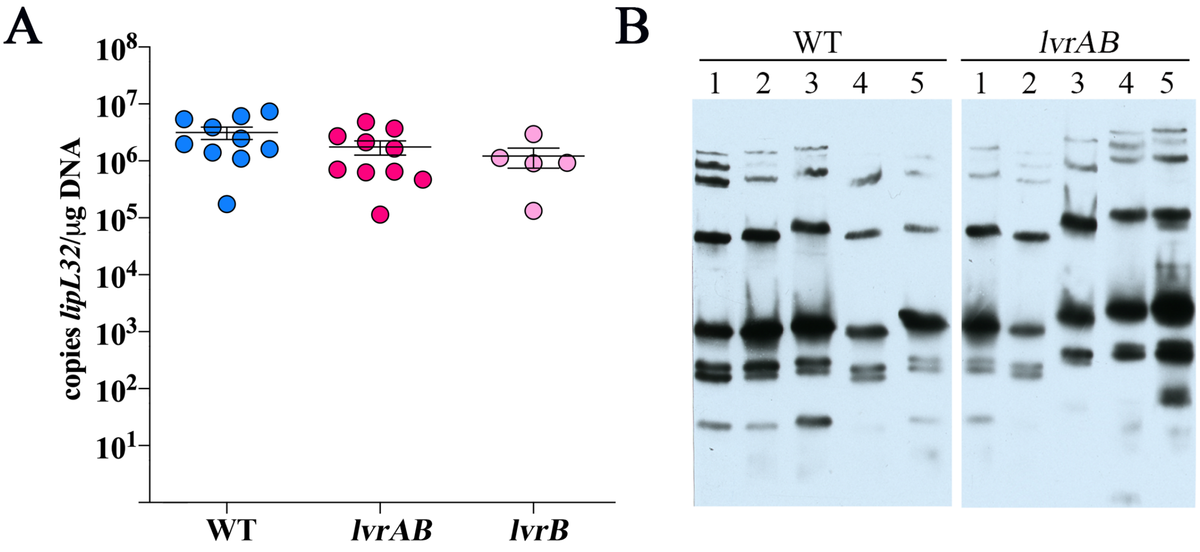

**S6 Figure. Expression of LvrAB requires at least one functional PerR homolog but the absence of LvrAB alone is not solely responsible for avirulence of the *perRA/B* double mutant. A.** Burdens of leptospires in kidneys harvested from mice in Figure 8B. DNA samples from kidneys harvested 28 days post-inoculation were assessed (in quadruplicate) by qPCR using a Taqman-based assay for *lipL32*. Bars represent the average and standard error of the mean. *p*-values were determined by comparing burdens in mice infected with wild-type (WT) and mutant strains at the same timepoint using a two-tailed *t*-test; we saw no significant difference ($p>0.05$) between burdens between the WT, *lvrAB* and *lvrB* strains. **B**. Immunoblot analysis of sera collected from C3H/HeJ mice 28-days following intraperitoneal inoculation with $10^5$ wild-type or *lvrAB* mutant strains and then used to probe whole cell lysates of *L. interrogans* sv. Manilae strain L495 grown in EMJH at 30°C.

**Table S1. Summary of RNA-Seq raw read data.**

| Sample | Total number of mapped reads | Number of uniquely mapped reads | Number of reads mapped to CDSs* |
|---|---|---|---|
| WT #1 | 18,087,357 | 17,505,315 | 12,360,862 |
| WT #2 | 18,362,489 | 17,375,505 | 9,799,876 |
| WT #3 | 14,949,092 | 14,258,122 | 8,343,892 |
| *perRA* #1 | 13,154,366 | 12,472,254 | 8,295,172 |
| *perRA* #2 | 14,682,864 | 14,133,388 | 9,695,747 |
| *perRA* #3 | 16,863,642 | 16,213,387 | 11,771,152 |
| *perRB* #1 | 18,394,538 | 17,552,727 | 8,701,072 |
| *perRB* #2 | 17,955,855 | 17,004,569 | 7,175,877 |
| *perRB* #3 | 17,788,952 | 16,702,526 | 8,054,671 |
| *perRA/B* #1 | 16,645,401 | 15,744,452 | 10,338,776 |
| *perRA/B* #2 | 17,448,069 | 16,770,321 | 11,931,747 |
| *perRA/B* #3 | 14,847,761 | 13,367,771 | 6,580,779 |

*after removing ncRNAs and pseudo-genes

**Table S2. Comparative RNA-Seq data for *L. interrogans* sv. Manilae L495 wild-type and *perRA* strains cultivated in dialysis membrane chambers (DMCs)**. The genome sequence of *L. interrogans* sv. Manilae strain UP-MMC-NIID LP (accession numbers NZ_CP011931.1, NZ_CP011932.1 and NZ_CP011933.1) was used for mapping and differential gene expression analysis. All non-coding RNAs and pseudogenes were removed before DESeq2 analysis. **Column A**: RefSeq locus tag. **Column B**: Locus tag. **Column C**: *L. interrogans* serovar Copenhageni strain Fiocruz L1-130 (accession numbers NC_005823.1 and NC_005824.1) orthologs identified using OrthoVenn 2.0 [3]. Dashes (-) indicate genes for which no clear ortholog was identified. **Column D**: Description of gene product, following genome annotation. **Columns E, F**: Clusters of Orthologous Group (COG) classifications based on MicroScope [4]. **Column G**: Fold-regulation of the corresponding gene based on RNA-Seq analysis by Zavala-Alvarado *et al.* [1, 2] performed using the same strains cultivated *in vitro*. Positive and negative numbers indicate upregulation and repression, respectively, in the WT compared to the mutant strain. Dashes (-) indicate genes that are not differentially regulated at least 3-fold ($p \leq 0.05$) by PerRA *in vitro*. **Column H**: Fold-regulation determined using DESeq2 based on WT vs. *perRA* mutant RNA-Seq analysis using leptospires cultivated within DMCs. **Column I**: Type of regulation by PerRA in DMCs. Genes expressed at ≥3-fold higher/lower levels in the WT vs. mutant with a False-discovery rate-adjusted-*p* value ($q$) ≤0.05 were considered differentially expressed. “NO” indicates genes that are not regulated by PerRA in DMCs; “Up” indicates genes upregulated by PerRA in DMCs (expressed at lower levels in the mutant vs. WT); “Down” indicates genes downregulated by PerRA in DMCs (expressed at higher levels in the mutant vs WT). **Columns J-O**: Number of mapped reads per gene for each one of the three biological replicates per strain. **Columns P-AC**: Output from DESeq2 for WT vs. *perRA* mutant strains cultivated in DMCs (3 biological replicates per strain). **Column P**: Mean DESeq2 values for each gene. **Column Q**: $\text{Log}_2$-fold change in gene expression. **Column R**: Power function transformation of $\log_2$-fold change. **Column S**: Fold regulation. **Column T-W**. Statistical analysis of differential gene expression including standard error estimate for the $\log_2$-fold change estimate (lfcSE, column T) and adjusted *p*-value (W). **Columns X-AC**: Normalized copy numbers per gene (3 biological replicates per strain).

**Table S5. Strain used in these studies.**

| Strain Name | Description | Antibiotics Resistance | Reference |
|---|---|---|---|
| WT | *L. interrogans* sv. Manilae st. L495, wild-type parent | - | [7] |
| *perRA* | *perRA*::$Km^R$, *Himar1* Tn insertion at position 62 bp in *LIMLP10155* | Kanamycin | [7] |
| *perRB* | *perRB*::$Km^R$, *Himar1* Tn insertion at position 287bp in *LIMLP05620* | Kanamycin | [2] |
| *perRA/B* | *perRB*::$Km^R$ Tn mutant containing an insertion within *perRA*, generated by allelic replacement | Kanamycin, Spectinomycin | [2] |
| *lvrB* | *lvrB*::$Km^R$, *Himar1* Tn insertion in *lvrB*/*LIMLP08485* | Kanamycin | [8] |
| *lvrAB* | *lvrAB*::$Km^R$, *Himar1* Tn mutant insertion in *lvrA*/*LIMLP08490* | Kanamycin | [8] |

**Table S6. Oligonucleotide primers used in these studies.**

| Primer | Sequence (5'-3') | Purpose | Reference |
|---|---|---|---|
| lipL32-F | TTGGATCCGTGTAGAAAGAATGTC | qPCR/qRT-PCR | [9] |
| lipL32-F | TCGTCCAATTTTTGAACTGGTTT | qPCR/qRT-PCR | [9] |
| lipL32 probe | [6FAM]-CCAAATCGCCAAAGCTGCGAAAGC-[BHQ1] | qPCR/qRT-PCR | [9] |
| PerRA-F | AACGCTTCTCGTGCCACTAT | qRT-PCR | This study |
| PerRA-R | CGCGTGATGATGATGGACTA | qRT-PCR | This study |
| PerRB-F | TTTCCTGTTTGGGAAAATCG | qRT-PCR | This study |
| PerRB-R | CCAGAAATTCAGGTGGGAGT | qRT-PCR | This study |
| LIMLP04825-F | ACAGTCTGCGGAAAAATCGT | qRT-PCR | This study |
| LIMLP04825-R | TGTTCGTTGCAAGTTCCGTA | qRT-PCR | This study |
| LIMLP18590-F | CATCTGATGGAAACGGGAAC | qRT-PCR | This study |
| LIMLP18590-R | CACGGTCGCATTGTTTACAG | qRT-PCR | This study |
| Tn5'out | CCGAAGTTCCTATACTTTCTAGAGAATAGGA | Sequencing | This study |
| Tn3'out | AAGCTTTAACTACAAGCTTTTTAGACATCTAATC | Sequencing | This study |
| PerRAseq-F | ATGAAGGATTCTTACGAAAGAAGC | Sequencing | This study |
| PerRAseq-R | TTATGGATTTTTTTGCCTTTGAGTGTAATG | Sequencing | This study |
| PerRBseq-F | ATGGAATCGTTATTTGCTAAAAAAGTTTGC | Sequencing | This study |
| PerRBseq-R | TTAAGTTTCTGAAACCAGATTTCCGGTAAG | Sequencing | This study |
| PerRAout-F | TCAAATTCTTTAAAATTAAAAATTTATTTC | Sequencing | This study |
| PerRAout-R | AAAACTTAGGTTTTTCTGTAAAATGATG | Sequencing | This study |
| FlaB1/pAE-F | TCACCTCGAGGGATCCATGATCATTAATCACAACT TAGCCG | Cloning | This study |
| FlaB1/pAE-R | TCTCGAGGTCGGATCCTTATCTAAGAAGCTGCAGA ACCG | Cloning | This study |
| PerRA/pET28a-F | CGCGCGGCAGCCATATGAAGGATTCTTACGAAAGA AGCAAAA | Cloning | This study |
| PerRA/pET28a-R | GTCATGCTAGCCATATGTTATGGATTTTTTTGCCT TTGAGT | Cloning | This study |
| PerRB/pET28a-F | CGCGCGGCAGCCATATGGAATCGTTATTTGCTAAA AAAGTTT | Cloning | This study |
| PerRB/pET28a-R | GTCATGCTAGCCATATGTTAAGTTTCTGAAACCAG ATTTCCG | Cloning | This study |
| LvrA/pET28a-F | CGCGCGGCAGCCATATGATAAATTCAACCGATTTA AAACACA | Cloning | This study |
| LvrA/pET28a-R | GTCATGCTAGCCATATGTTATATCGTTTCTAAAAAT GTCCT | Cloning | This study |
| LvrB/pET28a-F | CGCGCGGCAGCCATATGAATAAATGGAAATTCCTT TTCTTAG | Cloning | This study |
| LvrB/pET28a-R | GTCATGCTAGCCATATGTTATTGAACACTTGGGTCG | Cloning | This study |
